# DR5 CAR-T cells target solid tumors and suppress MDSCs with minimal toxicity

**DOI:** 10.1101/2025.06.09.658618

**Authors:** Huaishan Wang, Shujing Liu, Prithvi Sinha, Fang Liu, Xiaogang Zhang, Yeye Guo, Beatriz Goncalves, Qiuxiang Zheng, Haiwei Mou, Jingbo Yang, Lili Huang, John Scholler, Fei Miao, Tingting Zeng, Giorgos Karakousis, Alexander C. Huang, Tara Mitchell, Ravi Amaravadi, Lynn Schuchter, Michael Milone, Wei Guo, Carl June, Meenhard Herlyn, Yi Fan, Xiaowei Xu

## Abstract

Chimeric antigen receptor (CAR) T cell therapies have poor efficacy in solid tumors due to limited target specificity and an immunosuppressive tumor microenvironment. We investigated death receptor 5 (DR5) as a CAR target based on its high expression in both solid tumors and myeloid- derived suppressor cells (MDSCs). We engineered agonistic DR5-specific CAR constructs and evaluated their activity in multiple models, demonstrating DR5-expression-dependent tumor killing, confirmed by knockout and overexpression experiments. DR5-targeting single-chain variable fragments retained their pro-apoptotic activity when expressed on non-effector cells or extracellular vesicles. Among multiple CAR designs, we identified a construct with optimized binding affinity that maintained T cell viability while preserving strong tumor and MDSC-killing potency. To assess safety and efficacy in an immunocompetent setting, we also developed a murine DR5-targeted CAR. In multiple xenograft and syngeneic mouse models, DR5 CAR-T cells reduced tumor growth, prolonged survival, and did not cause detectable toxicity. In patient- derived organoids and tissue slices, DR5 CAR-T cells infiltrated tumor tissues, reduced MDSCs, boosted CD8+ T cell activity, and inhibited tumor growth. These findings support DR5-targeted CAR-T therapy as a promising strategy for treating solid tumors, which combines direct tumor cytotoxicity with immune activation and minimizes off-target effects.

****Teaser**:** DR5 CAR-T cells eliminate tumor cells and MDSCs and activate tumor-resident CD8+ T cells within the TME.

## Introduction

Chimeric antigen receptor (CAR) T cell therapies have shown remarkable success in hematologic malignancies but remain largely ineffective in solid tumors. This is due in part to the lack of tumor-specific antigens and the challenges posed by the immunosuppressive tumor microenvironment (TME) ^1–4^. The TME includes various cell types that suppress immune activity, such as myeloid-derived suppressor cells (MDSCs), cancer-associated fibroblasts (CAFs), and tumor-associated macrophages (TAMs) ^5–8^. These cells, along with dense extracellular matrix components, limit CAR-T cell infiltration and contribute to T cell exhaustion, ultimately reducing cytotoxic efficacy. Targeting suppressive elements of the TME, particularly MDSCs, is an emerging strategy to improve CAR-T function in solid tumors ^9,10^.

Death receptor 5 (DR5, also known as TRAIL-R2 or CD262) is a transmembrane receptor that plays a crucial role in apoptosis signaling. It is upregulated in many solid tumors, including melanoma, breast, lung, and colon cancers, and is also expressed on MDSCs ^11^. Activation of DR5 by TRAIL ligands or agonistic antibodies initiates a complex signaling cascade involving downstream effectors such as FADD, caspase-8, BID, and caspase-3 ^12,13^. This selective expression pattern has made DR5 an attractive therapeutic target ^13–16^. Although several DR5 agonist antibodies have been tested in clinical trials and shown to reduce MDSCs, their overall antitumor efficacy has been limited ^15–18^. MDSCs play a critical role in suppressing immune responses and promoting tumor progression, angiogenesis, metastasis, ^19^, and resistance to checkpoint inhibitors ^20^. Eliminating MDSCs may help reverse immunosuppression in the TME and improve the performance of both CAR-T cells and endogenous immune cells.

In this study, we engineered four DR5-targeting CAR constructs and evaluated their cytotoxicity in both 2D and 3D tumor models. DR5-targeting scFvs retained pro-apoptotic activity when expressed on non-effector cells or delivered via extracellular vesicles. Using affinity measurements and functional screening, we selected a construct that maintaining high affinity and potency, while minimized T cell fratricide. To evaluate safety and function in an immunocompetent setting, we also generated a murine DR5 CAR. DR5 CAR-T cells demonstrated antitumor activity in xenograft and syngeneic models without evidence of off-tumor toxicity. In patient-derived tumor models, these cells infiltrated tumors, reduced MDSCs, activated tumor-resident CD8+ T cells, and inhibited tumor growth. Together, these results support DR5-targeted CAR-T cells as a promising dual-function immunotherapy capable of inducing direct tumor cell death while reshaping the tumor immune landscape.

## Results

### DR5 CAR-T cells are highly cytotoxic to DR5^+^ tumor cells

To evaluate the efficacy of human DR5 CAR T cells, the protein sequences of four human DR5- scFvs were reverse-translated into DNA sequences, codon-optimized, and cloned into the pTRPE lentiviral vectors **(Fig. 1A, S1)**, and confirmed by Sanger sequencing. T-cells from freshly collected human peripheral blood mononuclear cells (PBMC) were expanded and transfected with lentiviruses carrying different DR5-scFv CAR constructs (1#-4#). After magnetic bead sorting, most of the T-cells expressed DR5-scFvs **(Fig. 1B)**. Melanoma (A375), ovarian cancer (OVCAR5), and hepatocellular carcinoma (HepG2) expressed similar levels of DR5 **(Fig. 1C)**. DR5 CAR-T cells were highly cytotoxic to these DR5^hi^ tumor cells **(Fig. 1D)**. Among the killing assays, DR5 CAR-T cells 1# and 3# showed more cytotoxicity to tumor cells than the other two DR5 CAR-T cells *in vitro*. To study the treatment efficacy of DR5 CAR-T cells against melanoma cells in 3D cultures, bicellular spheroids composed of GFP^+^ A375 cells and BJ fibroblasts were cocultured with DR5 CAR-T cells at E:T ratio of 1:1. Cells in the spheroids were stained with Hoechst33342 and GFP intensity was analyzed **(Fig. 1E-F)**, untransduced (UTD) T cells and CD19 CAR-T cells were included as controls. The results showed that DR5 CAR-T cells effectively killed DR5+ melanoma cells in 3D spheroids. To further evaluate the activation and function of DR5 CAR-T cells, we performed reverse-phase protein arrays (RPPA) to analyze CAR-T cell activation after coculture with A375 target cells **(Fig. S2A)**. Representative proteins involved in T-cell activation are shown in **Fig. 1G**. The data demonstrated that immune activation-associated proteins such as PDK1 and granzyme B levels were elevated upon incubation with the target cells, whereas UTD cells remained resting. Proteins associated with cell death and apoptosis were highly expressed in the target cells cocultured with DR5 CAR-T cells, compared to those cocultured with UTD and target cells alone **(Fig. S2B)**. These findings demonstrate that DR5 CAR T cells effectively kill DR5-expressing tumor cells in 2D and 3D models.

**Fig. 1.**
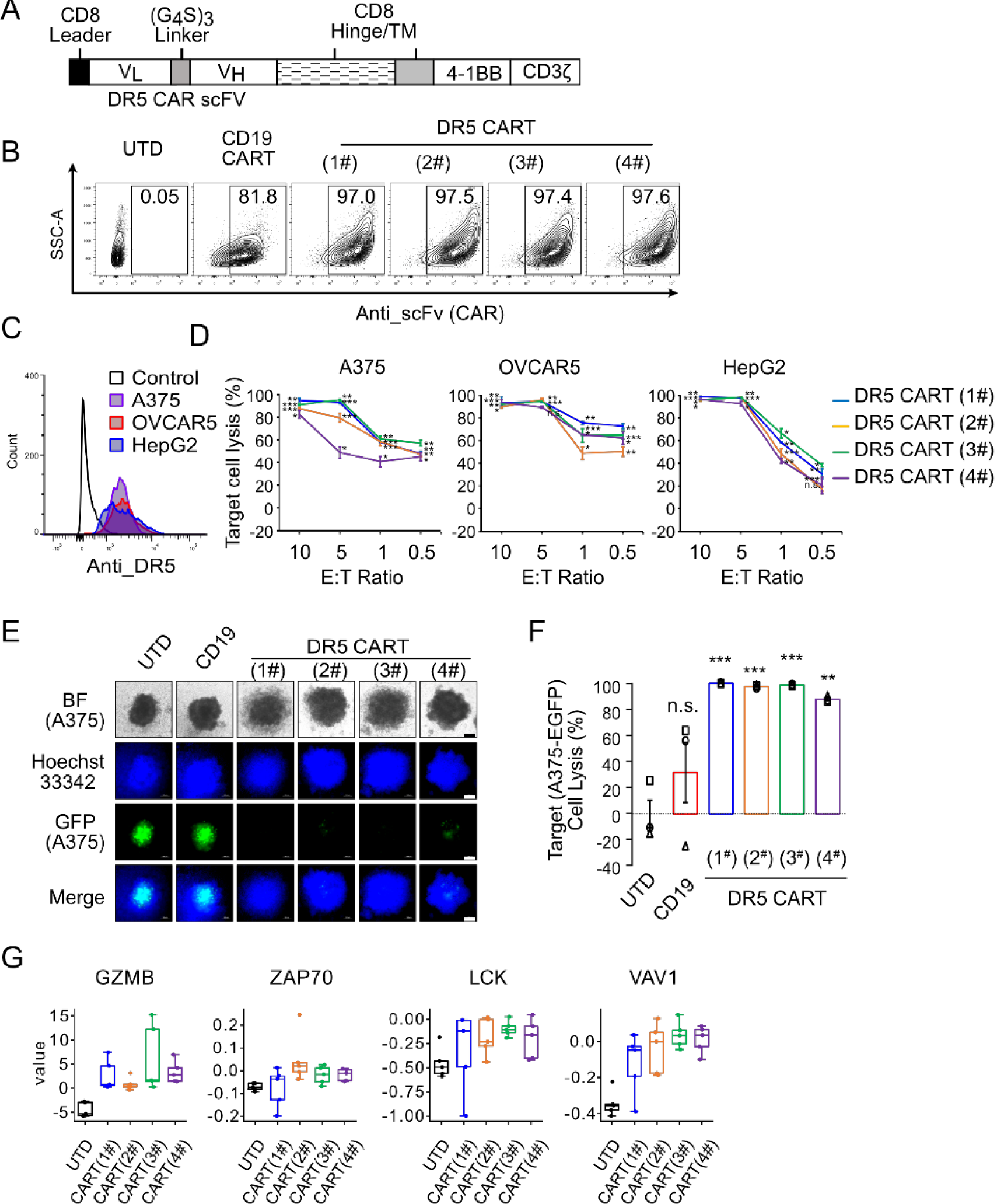
DR5 CAR-T cells are cytotoxic to DR5^+^ cancer cells. (A) Schematic of DR5 chimeric antigen receptors construct architecture. (B) Expression of the CAR on transduced T cells after magnetic bead purification. (C) DR5 expression on Luciferase- GFP transfected melanoma cell line A375, ovarian cancer cell line OVCAR5, and hepatocellular carcinoma cell line HepG2. (D) Luciferase-based cytotoxicity assay after 16 hours of coculture of DR5 CAR-T cells with cancer cells at indicated E:T ratio, n=3. (E) Representative images of 3D bicellular spheroids of A375 cells (GFP+Hoechst33342+) and fibroblast cell line BJ (Hoechst33342+) coculture with indicated DR5 CAR-T cells at E:T ratio of 1. Bar=100 μm. (F) Cytotoxicity of DR5 CAR-T to A375/BJ bicellular spheroids, target cell lysis data shown as loss of GFP signal density, n=3. (G) PRRA assay analysis of representative T cell activation gene expression after co-culture with A375 melanoma cells for 4 hours, n=5. All data shown as Mean±S.E.M., *P<0.05, **P<0.01, ***P<0.001.

### DR5 CAR-scFvs on cell and EV surfaces effectively induce target cell lysis

Since agonistic DR5 antibodies can directly activate the DR5 pathway and induce target cell apoptosis ^13,17^, we studied whether the DR5-scFvs can directly induce target cell death without the need for T cell activation. SupT1 cell is a T lymphoblast cell line that is not cytotoxic to other cells. We generated stable DR5 CAR-expressing SupT1 cells and confirmed the expression of DR5 CARs on the cell surface **(Fig. 2A)**. SupT1 cells were not cytotoxic and did not express cytotoxic T cell functional molecules, such as granzyme B, perforin and IFN-γ **(Fig. 2B)** or the target antigen DR5 **(Fig. S3A)**. We evaluated the cytotoxic effect of DR5 CAR-SupT1 cells on melanoma cells with various levels of DR5 expression **(Fig. S3B)**. DR5 CAR-SupT1 cells effectively killed these melanoma cells in a dose- and DR5 expression level-dependent manners **(Fig. 2C)**. We performed Hoechst33342/PI double staining assays and confirmed the tumor-killing by DR5 CAR-SupT1 cells **(Fig. 2D-E)**. We also tested the effect of DR5-CAR SupT1 cells on non-adherent lymphoblast cell K562-CD19 cells, which express a lower level of DR5 than A375 **(Fig. S3A)**, DR5-CAR SupT1 cells induced K562-CD19 cell apoptosis and death **(Fig. S3C-D)**. We also transfected SupT1 cells with CD19-CAR. In contrast, the CD19-CAR SupT1 cells did not show similar cytotoxic effects to K562-CD19 cells **(Fig. S3C-D)**. These data support that membrane DR5 scFvs induce DR5 activation in target cells.

**Fig. 2.**
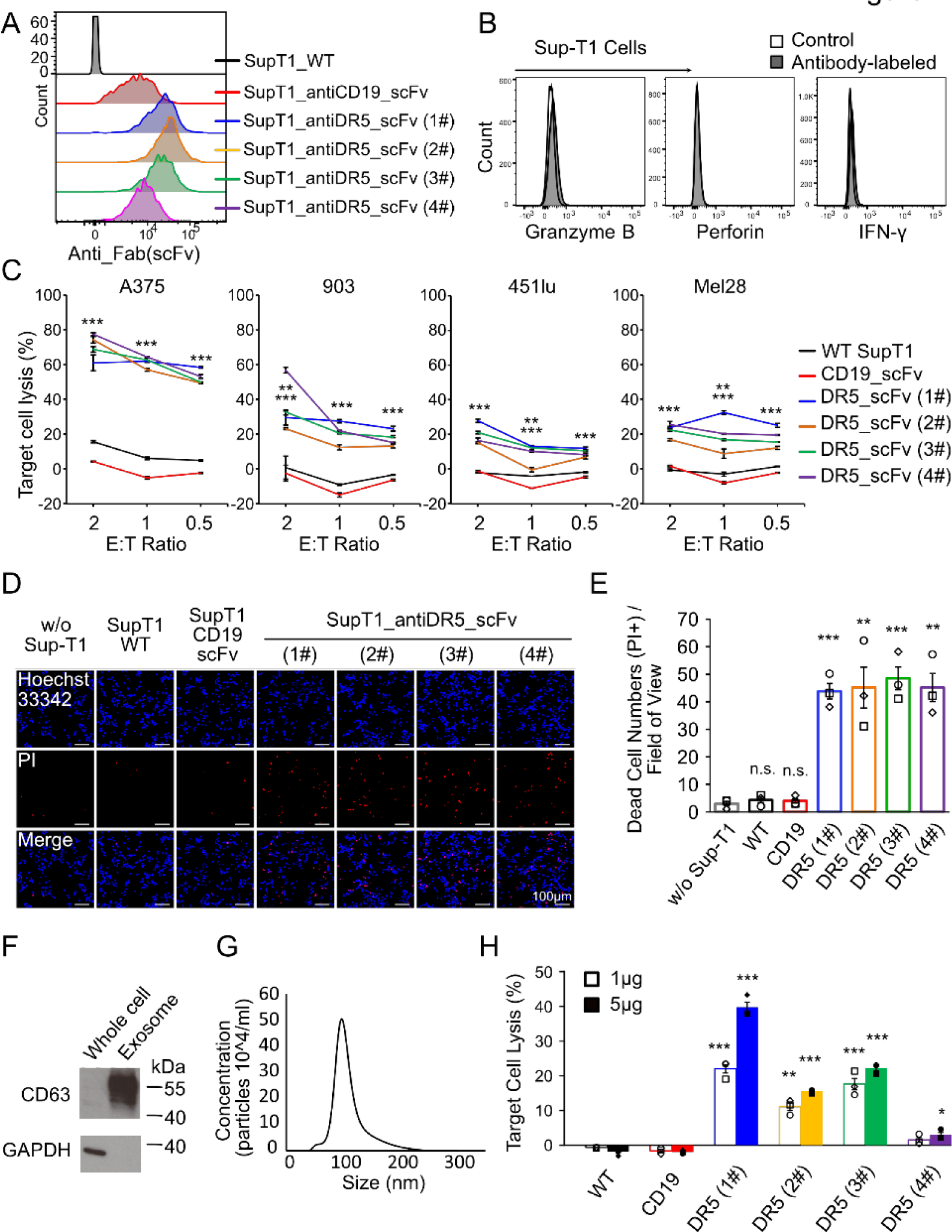
Membrane DR5-scFvs induce target cell death. (A) DR5-scFv membrane expression on non-cytotoxic SupT1 cells. SupT1 cells are transfected with CD19 and DR5 CAR constructs. CAR expression on the cell surface is detected using FACS. (B) SupT1 cells do not express cytolytic proteins. Cytotoxic T cell functional protein expression in SupT1 cells by FACS. (C) Cytotoxicity of DR5 CAR-SupT1 cells against melanoma cells. LDH- releasing assays are performed 16 hours after co-culture of DR5 CAR-SupT1 and melanoma cells at indicated E:T ratios, n=3. (D) Representative images of Hoechst33342/PI double staining assays showing DR5 CAR-SupT1 cells kill A375 cells in 2D culture at an E:T ratio of 1:1, n=3. CD19 CAR-SupT1 and wild-type SupT1 cells were used as controls. (E) Statistical analysis of cytotoxicity of DR5 SupT1 to A375 cells. The number of PI+ (Red) cells in different fields was quantified and averaged, n=3. (F) Western blot analysis showing the sEV marker CD63 expression in SupT1-derived sEVs. (G) Size analysis indicated the sEV size at ∼100nm. EV size is measured by the NanoSight NS300 instrument. (H) Cytotoxicity of DR5 CAR-SupT1 sEVs. LDH releasing assay after coculture of sEVs from CAR-SupT1 cells and A375 melanoma at the indicated amount for 16 hrs, n=3. All data are shown as Mean±S.E.M., *P<0.05, **P<0.01, ***P<0.001 by Student’s t-test, n.s., no statistically significant difference.

**Fig. 3.**
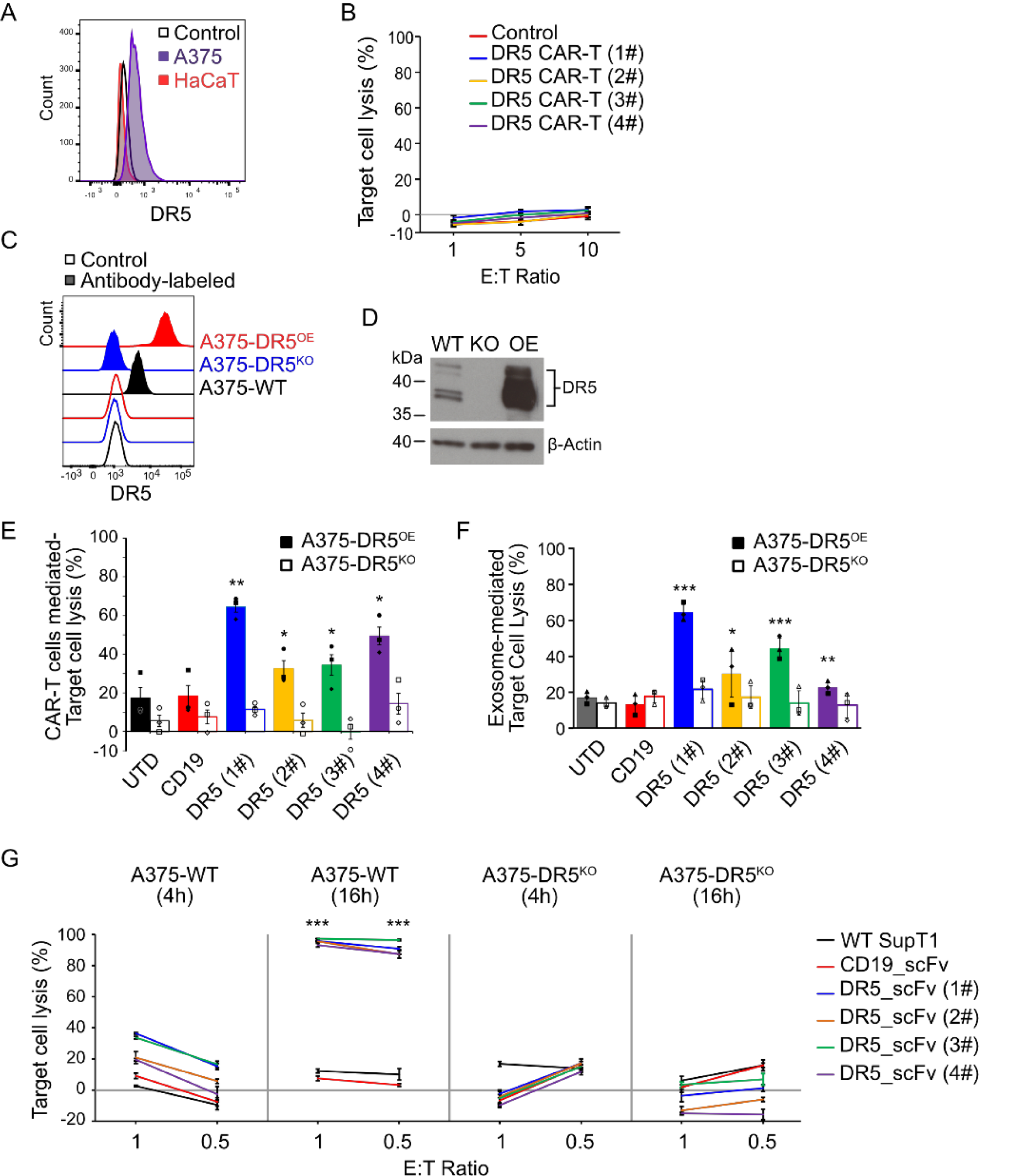
DR5 CAR-T cells are not cytotoxic to DR5-negative cells. (A) DR5 expression levels on different cell lines. HaCaT cells expressed a lower level of DR5 than A375 cells. (B) DR5 CAR-T cells with little toxicity to HaCaT cells at indicated E:T ratios, n=3. (C) Membrane DR5 expression levels in DR5 knockout (KO) and over-expression (OE) A375 cells. FACS was used to measure DR5 expression in these cell lines. (D) DR5 expression levels in DR5^KO^ and DR5^OE^ A375 cells. Western blot was used to measure DR5 expression in these cell lines. (E) Cytotoxicity assay of DR5 CAR-T cells to A375 cells. Luciferase-based assays were performed after coculture of DR5 CAR-T cells with different A375 cells for 16 hrs at the indicated E:T ratio of 5:1, n=3. (F) Cytotoxicity assay of sEVs derived from DR5 CAR-T cells to A375 cells. Luciferase-based assays are performed after coculture of sEVs (1 μg/ml) from DR5 CAR-T cells with different A375 cells for 16 hrs, n=3. (G) Luciferase assays after 4 or 16 hours of DR5 CAR SupT1-A375 coculture at indicated effector to target (E:T) ratios to measure cytotoxicity by DR5 CAR-SupT1 cells against DR5 wild type and knockout A375 cells, n=3. All data shown as Mean±S.E.M., *P<0.05, **P<0.01, ***P<0.0.

Extracellular vesicles (EVs) are nanosized lipid bilayer particles that are naturally released from almost all types of cells ^21^. Based on size and isolation methods, EVs are further divided into small extracellular vesicles (sEVs) and microvesicles (MVs) ^22^. We investigated the cytotoxicity of sEVs and MVs derived from DR5-CAR-SupT1 cells on melanoma cells. sEVs were enriched in CD63 and had a size of ∼100 nm **(Fig. 2F-G).** These sEVs effectively induced A375 melanoma cell death (**Fig. 2H**). Microvesicles (MV) (>200nm) from DR5-CAR-SupT1 cells also had similar effects **(Fig. S4)**. These findings support that membrane DR5 CAR-scFvs can induce target cell death by directly activating DR5 pathways, independent of T-cell activation.

### DR5 CAR-T cells are not cytotoxic to DR5^-^ cells

To evaluate the specificity and off-target toxicity of DR5 CAR-T cells, we examined their cytotoxicity to DR5^-^ HaCaT keratinocyte cells **(Fig. 4A)**. The results demonstrated that all four DR5 CAR-T cells were not cytotoxic to HaCaT cells **(Fig. 4B)**. We then generated DR5 overexpression (OE) or knockout (KO) A375 melanoma cells. Flow cytometry and western blot confirmed the expression of DR5 in these cells, respectively (**Fig. 4C-D)**. Luciferase-releasing assays demonstrated that neither DR5 CAR-T cells nor EVs derived from CAR-T cells exhibited significant toxicity to DR5^KO^ cells **(Fig. 4E-F).** Furthermore, surface-anchored DR5-scFvs on SupT1 cells demonstrated no significant cytotoxic effect on DR5^KO^ A375 cells **(Fig. 4G)**. These results support that DR5 CAR-T cells are target-specific and do not exhibit off-target toxicity to DR5^-^ cells.

**Fig. 4.**
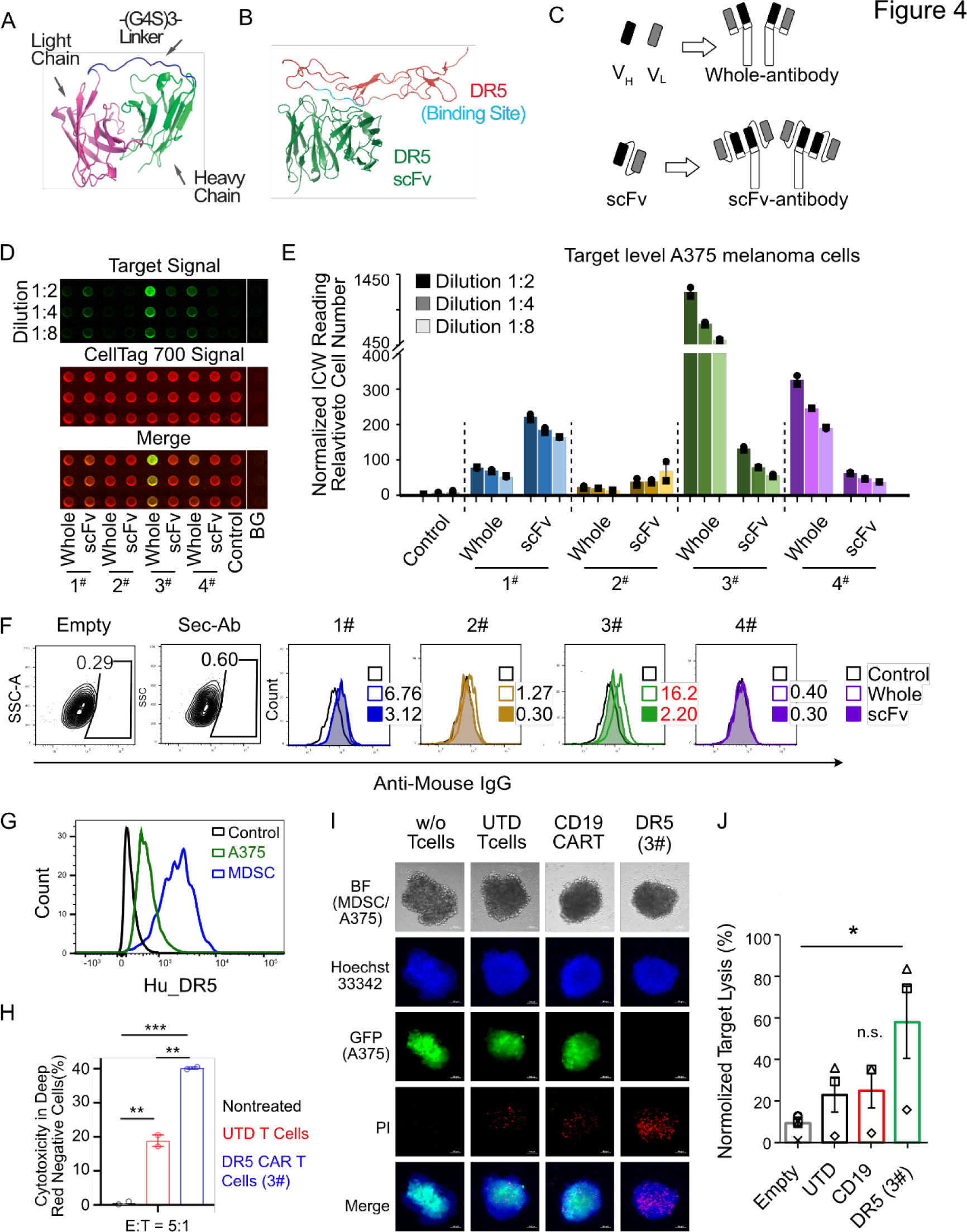
Binding affinity of DR5-scFv to its target. (A) Representative scFv structure of anti-DR5 (1#) with light chain region (pink), heavy chain region (green), and G4S linker (blue). (B) Representative binding of anti-DR5 scFv (Green) to DR5 protein (Deep Red). (C) Schematics of DR5 whole antibodies (upper panel) and DR5 scFv- antibodies (lower panel). (D) Image panels of On-Cell Western (OCW) assay showing DR5 antibody binding intensity at the indicated dilutions. Pre-fixed A375 cells were used in the binding assays. (E) Normalized OCW data of DR5 antibody binding intensity to A375 cells, shown as Mean±S.E.M. of 3 wells. (F) Representative flow cytometry data showing the DR5 antibody binding to A375 cells. Different DR5 whole and scFv antibodies (4μg) were used in the assays. (G) DR5 expression on the surface of MDSCs and A375 melanoma cells. (H) Cytotoxicity of DR5 CAR-T cells against MDSCs. LDH-releasing assays are performed after co-culture of T cells-target cells at E:T ratio 5:1 for 16 hours, n=3. (I) Representative images of 3D spheroids with A375 cells (GFP+Hoechst33342+) and MDSCs (Hoechst33342+) coculture with indicated DR5 CAR-T cells at E:T ratio of 1:1. Bar=100μm. (J) Statistical analysis of cytotoxicity of DR5 CAR-T cells against A375-MDSC spheroids, target cell lysis data shown as PI signal density, n=3-5. All data are shown as Mean±S.E.M., *P<0.05, **P<0.01 by Student’s t-test, n.s., no statistically significant difference.

### DR5-scFv affinity to DR5 correlates with its target-killing efficacy

CAR affinity to its target is crucial for CAR-T cell function and CAR-T cell-induced adverse reactions ^23–25^. To select the best CAR for further development, we first analyzed the affinity of all four DR5 scFvs to DR5 using in silico computational methods **(Fig. 4A and B).** The DR5 scFvs appear to bind to slightly different areas of DR5, however, the HADDOCK Scores of DR5- scFvs to DR5 binding affinity were not statistically **(Fig. S5A-B)**. To experimentally study the binding affinity, we constructed four different DR5 whole-antibodies and their corresponding DR5-scFv-antibodies **(Fig. 4C)**. Then, we evaluated the binding of these DR5 antibodies to pre- fixed A375 cells using On-Cell Western assays **(Fig. 4D)**. The normalized data of DR5 whole- antibody and DR5-scFv-antibody-binding to A375 cells are shown in **Fig. 4E**. DR5 antibodies bound to their target with different affinities, and the clone 1# DR5-scFv-antibodies had the highest affinity to their targets, followed by clone 3#. Flow cytometry data demonstrated that DR5 whole-antibodies and DR5-scFv-antibodies bound to A375 cells, similar to the results using On-Cell Western assay (**Fig. 4F, S6**). We then tested the effects of CAR expression on T cell proliferation. The viability of cultured DR5 CAR-T cells was measured, and clone 1# had the least viable cells (**Fig. S7A**). Similar results were observed after DR5 CAR-T cells were cocultured with A375 cells for 24 hours (**Fig. S7B**). Therefore, clone 3# CAR was selected for further development.

### DR5 CAR-T cells are cytotoxic to MDSCs *in vitro*

DR5 is highly expressed in MDSCs ^11^. To investigate the effects of DR5 CAR-T cells on MDSCs, we first generated MDSCs from PBMC-monocytes and confirmed their phenotype by flow cytometry, which showed typical CD11b+CD33+PDL1^hi^ expression **(Fig. S8A)**. We then co- cultured DR5 CAR-T cells with MDSCs and assessed T cell proliferation using CFSE staining. MDSCs significantly suppressed T cell proliferation compared to T cells alone **(Fig. S8B)**. Next, we evaluated DR5 expression on the surface of MDSCs using flow cytometry. MDSCs expressed a higher level of DR5 than the A375 melanoma cells **(Fig. 4G).** To evaluate the cytotoxicity of DR5 CAR-T cells against MDSCs, we performed LDH-releasing assays after 16 hours of CAR- T-cells and MDSC coculture. The results showed that selected 3# DR5 CAR-T cells were significantly cytotoxic to MDSCs *in vitro* **(Fig. S4H)**. We then studied the treatment efficacy of DR5 CAR-T cells against 3D spheroids composed of GFP^+^ A375 cells and MDSCs at an E:T ratio of 1:1. The spheroids were stained with Hoechst33342 and PI and imaged using a fluorescent microscope **(Fig. 4I)**. The results showed that the DR5 CAR-T cells effectively induced lysis of target A375 cells in the spheroids based on overall GFP and PI staining **(Fig. 4I-J).** UTD-T cells or CD19 CAR-T cells were included as controls. These results demonstrate that DR5 CAR-T cells are cytotoxic to tumor cells in the presence of MDSCs.

### Human DR5 CAR-T cells inhibit melanoma growth *in vivo* in xenograft models

To investigate the effect of DR5 CAR-T cells *in vivo*, we first established an A375 melanoma xenograft model by injecting 5×10^6^ A375-DR5^OE^ with GFP and FFLuc melanoma cells or A375 wild type (WT) cells into the flanks of nude mice. The treatment schema is shown in **Fig. 5A**. Mice were treated with intratumoral injection of UTD T cells or DR5 CAR-T cells. Tumor bioluminescence images and Day 20 bright field tumor images are shown in **Fig. 5B**. The tumor bioluminescence kinetics of A375-DR5^OE^ tumor growth in the xenograft model are shown in **Fig. 5C**, indicating a significant reduction of tumor burden after DR5 CAR-T cells treatment compared to the UTD T cell control group. Moreover, the treated mice exhibited no significant body weight loss, indicating no apparent systemic toxicity **(Fig. 5D)**. All the mice were sacrificed, and histological analysis of the mouse tissues collected on day 20 revealed the absence of pathological changes in the major organs. To further confirm the effect of DR5 CAR-T cells *in vivo*, WT A375 melanoma cells were treated with intratumoral UTD T cells or DR5 CAR-T cells. Tumor growth was significantly reduced, and mouse survival was prolonged considerably (**Fig. 5E-G**). Body weight loss or systemic toxicity was not observed in the treated mice by histology. These results support that DR5 CAR-T cells can control melanoma tumor growth *in vivo*.

**Fig. 5.**
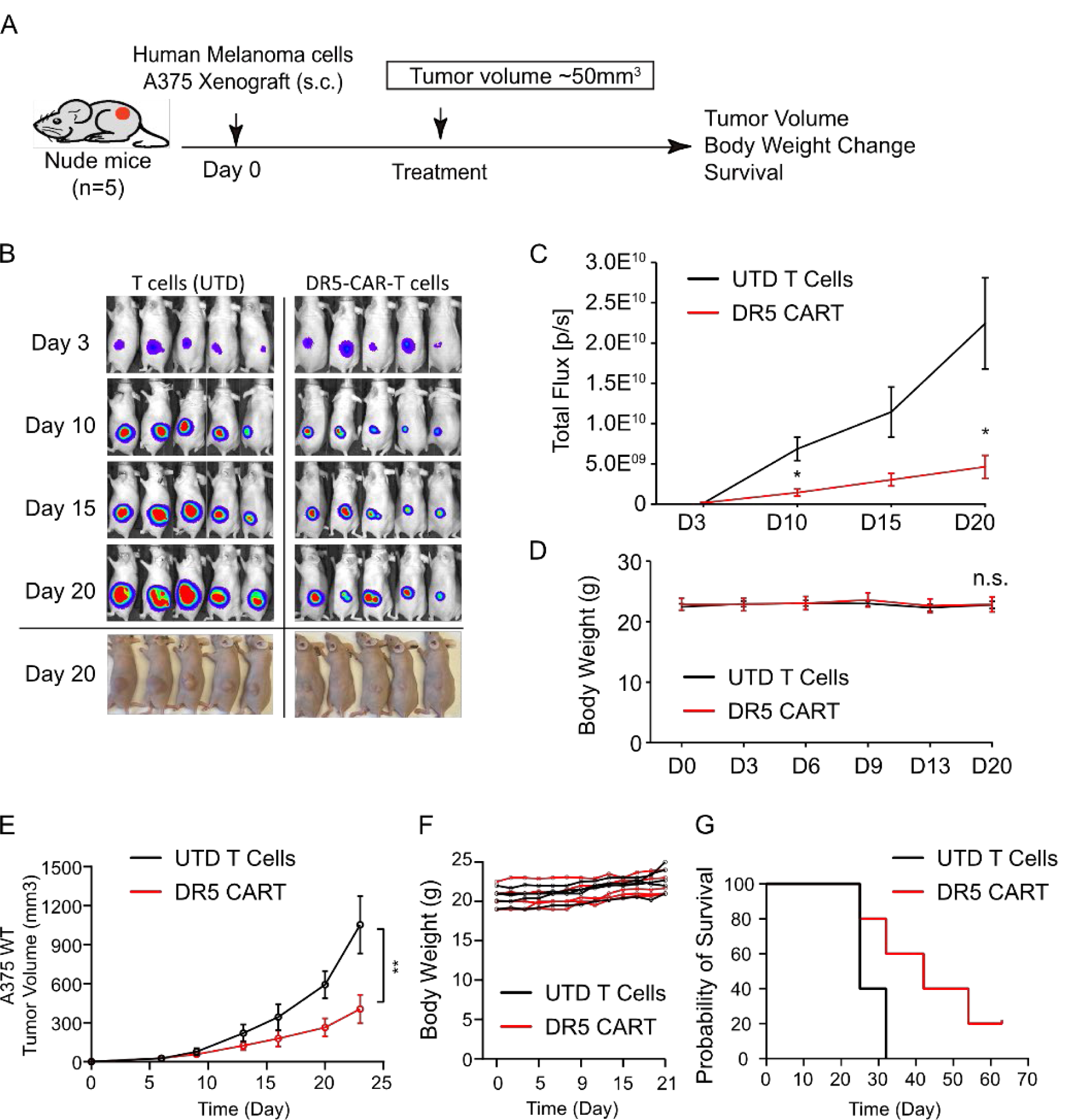
Effects of DR5 CAR-T cells *in vivo*. (A) Treatment schema of intratumoral DR5 CAR-T cells in A375 melanoma xenograft model. 5×10^6^ A375-DR5^OE^ (B-D) or A375 wildtype (WT) (E-G) melanoma cells are intratumorally injected into the flanks of nude mice, 5x10^6^ CAR+ T cells and same number of UTD cells in 100ul PBS were injected. (B) Representative A375-DR5^OE^ tumor bioluminescence images and Day 20 bright field tumor images. (C) Tumor bioluminescence kinetics of A375-DR5-GFP-FFLuc tumor growth in the xenograft nude mice model shown in (B), n=5. (D) Body weight curves of tumor- burden mice, n=5. (E) Tumor volumes of A375 WT tumor growth in the xenograft nude mice model, n=5. (F) Body weight curves of tumor-burden mice of E, n=5. (G) Survival curve of A375 WT xenograft mice, n=5. All data are shown as mean±SEM. *P<0.05, **P<0.01, ***P<0.001, n.s., no statistically significant difference.

### DR5 CAR-T cells are efficacious and safe in the syngeneic melanoma models

Since DR5 may be expressed in some normal tissues, we designed a CAR against murine DR5. Murine CAR-T cells allow us to evaluate the safety and efficacy in an immunocompetent setting. The CAR is designed based on the M5-1 monoclonal antibody ^26^ and the construct has traceable BFP and nLuc as we previously described ^27^. Murine DR5 CAR-T cells were manufactured using mouse splenocytes **(Fig. 6A),** and 31.4% of the CAR-T cells showed BFP expression **(Fig. S9)**. When these murine DR5 CAR cells were injected into healthy C57 mice via the tail veins, these mice maintained their body **(Fig. S10)** and none died during treatment.

**Fig. 6.**
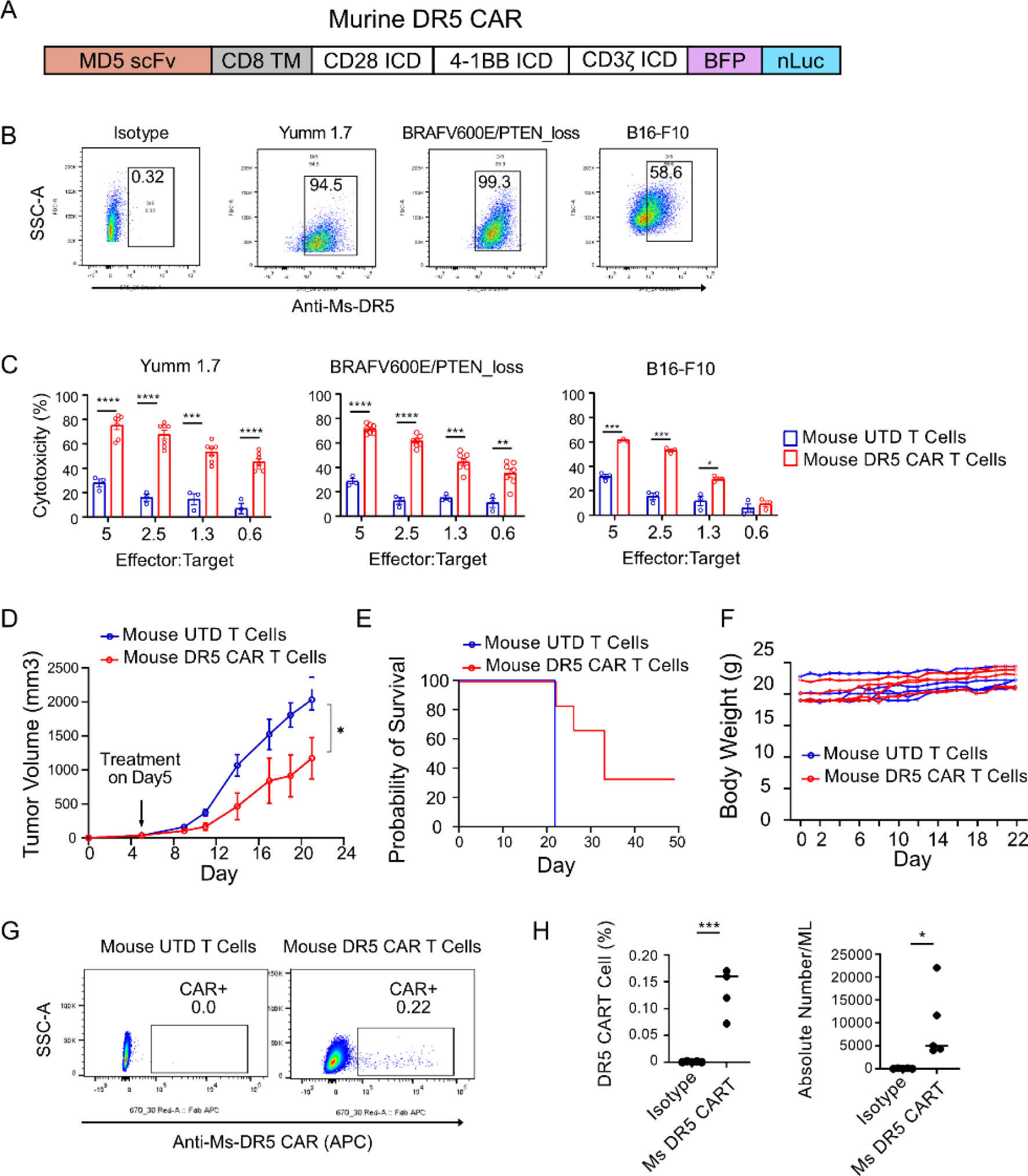
Murine DR5 CAR T cells are efficacious in syngeneic mouse models without on- target off-tumor toxicity (A) Schematic of murine DR5 chimeric antigen receptors construct architecture. (B) DR5 expression on mouse melanoma cell lines, Yumm 1.7, *BRAF^V600E^PTEN^-/-^*, and B16-F10, respectively. (C) Cytotoxicity assay after 12 hours of coculture of murine DR5 CAR-T cells with melanoma cells at the indicated E:T ratio, n=3. (D) Tumor volumes of Yumm 1.7 in syngeneic melanoma model treated with 5x10^6^ murine DR5 CAR T cells or murine UTD T cells via tail vein injection, n=5. (E) Survival curve of Yumm 1.7 syngeneic mice after treatment, n=5. (F) Body weight curves of tumor-bearing mice after treatment, n=5. (G) Representative flow cytometry data showing the circulating murine DR5 CAR T cells after sacrifice. (H) Murine DR5 CAR+ proportion of total mouse T cells and absolute number of DR5 CAR+ T cells. All data are shown as mean±SEM. *P<0.05, **P<0.01, ***P<0.001, n.s., no statistically significant difference.

We examined DR5 expression in several mouse melanoma cell lines and found that over 90% of Yumm1.7 and *BRAF^V600E^pTEN^-/-^* cells expressed DR5 and 59% of B16-F10 cells expressed DR5 **(Fig. 6B)**. Murine DR5 CAR-T cells induced significantly cytotoxicity in Yumm1.7 and *BRAF^V600E^pTEN^-/-^* cells even at E:T ratio of 0.6:1 after coculturing 12 hours **(Fig. 6C)** and 24 hours **(Fig S11)**. To further evaluate toxicity and the *in vivo* efficacy, Yumm 1.7 syngeneic melanomas were established in C57 mice. When the tumors became palpable, murine DR5 CAR cells were injected once through the tail veins. Systemic CAR-T cells led to significant tumor growth reduction and prolonged survival compared to UTD treated mice **(Fig. 6D-F)**. No weight loss **(Fig. 6F)** or histological changes were detected in the major organs. Flow cytometry confirmed the presence of circulating CAR-T cells after sacrificing the mice at the end of the experiment, indicating persistence of CAR-T cells following infusion **(Fig. 6G-H)**. These findings support the safety and therapeutic potential of DR5-targeted CAR-T cells in an intact immune system.

### DR5 CAR-T cells infiltrate and kill tumor cells in melanoma patient-derived organoids (MPDOs) and melanoma slice cultures (MSCs)

To evaluate the DR5 CAR-T cell efficacy against melanoma in patient tissues, we obtained fresh viable human melanoma tissues from surgically resected specimens, and the tumor tissues were sectioned into small fragments and cultured to form MPDOs. MPDOs were cultured with a mixture of labeled UTD T cells (pink, CellTrace Far Red) and DR5 CAR-T cells (green, CFSE). PI was used to demonstrate cell death. The live cell imaging studies demonstrated that few dead cells (red, PI+) were present in the MPDOs at the beginning of the experiments. Killing the cells in the organoids was relatively fast, and by 120 minutes, many dead cells in the MPDOs were stained positive for PI **(Fig. 7A and Movie S1).** Confocal images of MPDOs cocultured with DR5 CAR-T cells for 24 hours demonstrated that more infiltrating DR5 CAR-T cells and PI-positive dead cells were present in the MPDOs than MPDOs treated with UTD T-cells **(Fig. 7B).** CFSE- labeled DR5-CAR-T cells penetrated deep into the organoids, while UTD T-cells were mostly attached to the surface of the organoids. Imaging analysis was performed to quantify the fluorescence density in MPDOs and confirmed observation **(Fig. 7C).** After we cocultured the MPDOs with DR5 CAR-T cells for 48 hours, the organoids disintegrated as the organoids lost sharp tissue edges; the disintegration was not observed in the untreated MPDOs or MPDOs treated with UTD T-cells **(Fig. S12)**.

**Fig. 7.**
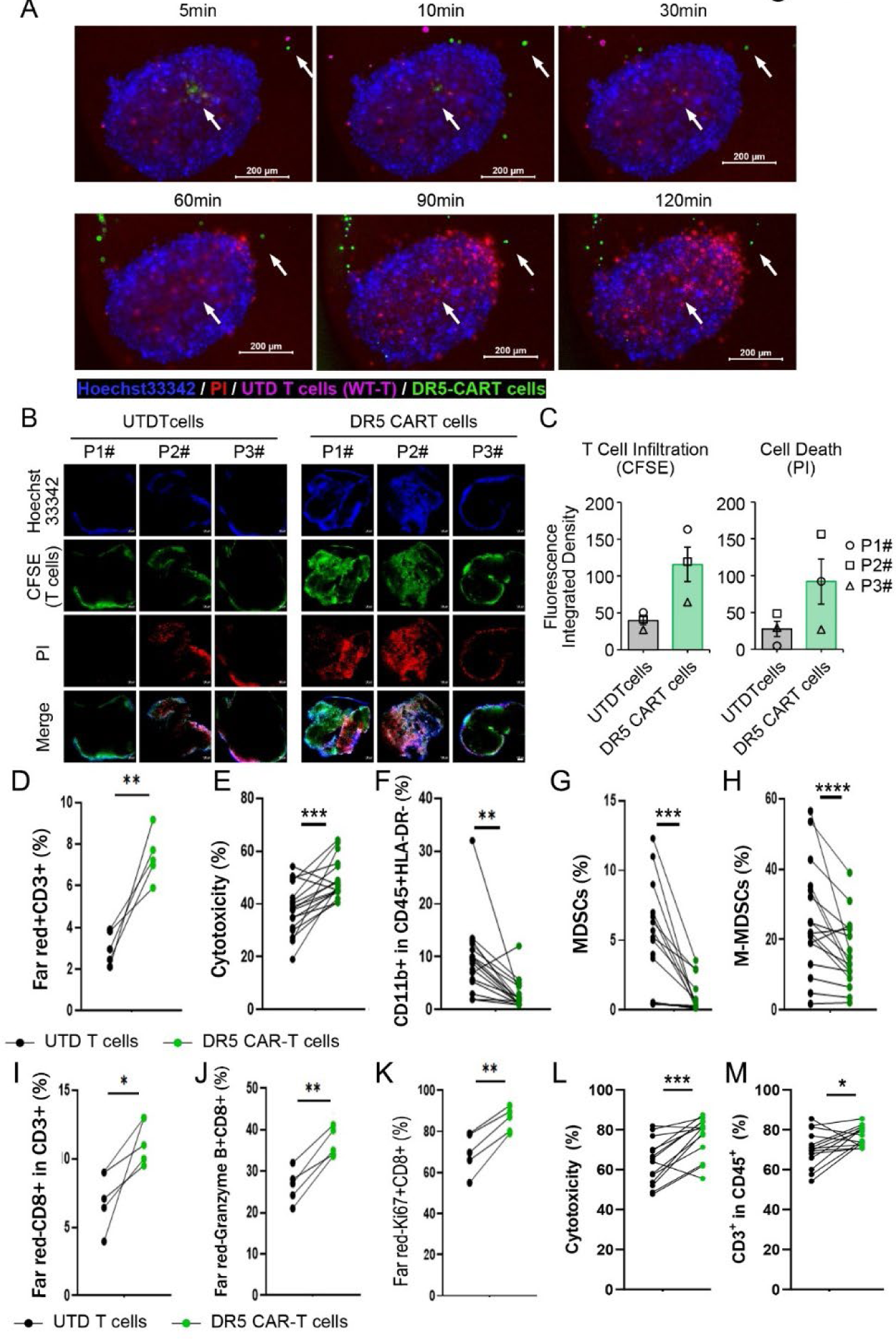
Effects of DR5 CAR-T cells on MPDOs and MSCs. (A) Representative fluorescence images of MPDOs co-cultured with UTD T cells (Pink) and DR5 CAR-T cells (Green). Arrows indicate the DR5 CAR-T cells, and arrowheads indicate dead tumor cells. Bars indicate 200 µm. Hoechst33342 and PI are used to label nuclei and dead cells, respectively. (B) Confocal images of MPDOs cocultured with DR5 CAR-T cells (CFSE) for 24 hours. Hoechst33342 and PI label nuclei and dead cells, respectively, n=3. (C) DR5 CAR-T cell infiltration and tumor killing in MDPOs. CAR-T cells are labeled with CFSE. Hoechst33342 and PI were used to label nuclei and dead cells, respectively. Fluorescent density is digitally analyzed, n=3. (D) DR5 CAR-T cells infiltrating MPDOs. UTD and CAR-T cells were labeled with far-red fluorescent dye. MPDOs from three melanoma patients were incubated with labeled UTD or CAR-T cells for 48 hours. After treatment, the MPDOs were collected and disassociated into single cells for FACS analysis. Far-red+CD3+ T cells are quantified. Each dot represents an independent well. (E-H) Effects of DR5 CAR-T cells on MDSCs. MPDOs from four melanoma patients were incubated with labeled UTD or CAR-T cells for 48 hours. After treatment, the MPDOs were collected and disassociated into single cells for FACS analysis. More cell death in the MPDOs was observed after CAR-T cell treatment (E). CD11b+CD45+HLA-DR- myeloid cells (F), CD33+CD11b+ MDSCs (G), CD14+CD1b+CD33+ M-MDSCs (H) were significantly decreased after CAR-T cell treatment, n=17, and each dot represents an independent well. (I-K) Tumor-resident CD8+ T cells were activated after CAR-T cell treatment. UTD and CAR-T cells were labeled with far-red fluorescent dye. MPDOs from three melanoma patients were incubated with labeled UTD or CAR-T cells for 48 hours. After treatment, the MPDOs were collected and dissociated into single cells for FACS analysis. Far-red-CD8+ T cells (I), Granzyme B+far-red- CD8+ T cells (J), Ki67+far-red-CD8+ T cells (K) were significantly increased after CAR-T cell treatment. n=5, and each dot represents an independent well. (L-M) Effects of DR5 CAR-T cells on MSCs. MSCs from five different patients were used in the experiment. One to two slices per patient were included in the study. Each dot represents a tumor slice. The MSCs were incubated with DR5 CAR-T cells or UTD cells for 48 hours. The tissues were washed and digested into single cells for FACS analysis. Cytotoxicity (L) and CD3+ T cells (M) are significantly increased after CAR-T cell treatment. All data are shown as Mean±S.E.M., *P<0.05, **P<0.01, ***P<0.001, ****P<0.0001.

MPDOs embedded in the Matrigel were treated with far-red-labeled DR5-CAR-T cells or UTD T cells for 48 hours. MPDOs were collected and dissociated into a single-cell suspension. FACS analysis showed that 3-fold more far-red+CD3+ CAR-T cells than far red+CD3+ UTD T cells were detected in the MPDOs **(Fig. 7D)**, consistent with the imaging analysis data **(Fig. 7C)**. Significantly more dead cells were present in the MPDOs after DR5 CAR-T cell treatment **(Fig. 7E)**. CD11b+ myeloid cells **(Fig. 7F)**, CD11b+CD33+ MDSCs **(Fig 7G)**, CD14+CD11b+CD33+ M-MDSCs **(Fig. 7H)**, and CD15+CD11b+CD33+ PMN-MDSCs **(Fig. S13)** were significantly decreased after DR5 CAR-T cell treatment compared to UTD T cell treatment. On the contrary, tumor-resident far-red-CD8+ T cells **(Fig. 7I)** were significantly increased, and these far-red- CD8+ T cells expressed significantly more granzyme B **(Fig. 7J)** and Ki67 **(Fig. 7K)**, indicating activation of tumor-resident CD8+ T cells after DR5 CAR-T cell treatment. To further confirm the effect of DR5 CAR-T cells, we also tested their effects in MSCs. MSCs represent a physiologically relevant culture system that preserves the original TME ^28–31^, demonstrating utility in predicting responses to small molecules ^30^ and checkpoint inhibitors ^32^. 300 μm thick fresh melanoma slices from 5 patients were incubated with UTD T-cells and DR5 CAR-T cells (2x10^6^) for 48 hours. DR5 CAR-T cells significantly increased the PI+ dead cells compared to MSCs treated with UTD T cells (**Fig. 7L**). MSCs were washed and lysed to form a single-cell suspension for FACS analysis. MSCs incubated with DR5 CAR-T cells had a significantly higher number of CD3+ T cells in the tissue than tissues treated with UTD T cells, supporting more tumor- infiltrating DR5 CAR-T cells in the tissues (**Fig. 7M**). These data suggest that DR5 CAR-T cells can infiltrate patient-derived melanoma tissues, inhibit the MDSCs, and activate tumor-resident CD8+ T cells.

## Discussion

CAR-T cell therapies face major barriers in solid tumors, particularly due to the immunosuppressive tumor microenvironment (TME) and the lack of tumor-specific antigens ^3^. In this study, we show that DR5-targeted CAR-T cells can overcome these limitations through a dual mechanism: directly killing DR5-expressing tumor cells and eliminating immunosuppressive MDSCs. These CAR-T cells exhibit strong antitumor activity *in vitro* and *in vivo*, including multiple xenograft and syngeneic mouse models, as well as patient-derived models, without evident toxicity to normal tissues. These CAR-T cells are capable of persisting in circulation. Therefore, DR5 is a promising CAR-T cell target that allows simultaneous inhibition of tumor cells and MDSCs.

One distinguishing feature of DR5 CAR-T cells is their ability to kill targets through both classical T-cell cytotoxic mechanisms and agonistic signaling via membrane-bound scFvs. We demonstrated that even non-cytotoxic Sup-T1 cells expressing DR5-scFvs could induce apoptosis in DR5+ tumor cells, and EVs derived from CAR-expressing non-cytotoxic cells retained this function. We recently demonstrated that DR5-engineered EVs released by natural killer cells were highly cytotoxic to tumor cells both *in vitro* and *in vivo* ^33^. These findings suggest that DR5 CAR-T cells have multiple ways to engage and destroy tumor cells, potentially improving efficacy in settings where conventional CAR signaling alone may not be sufficient.

DR5 is an appealing target due to its high expression in many solid tumors and MDSCs, with limited expression in normal tissues ^14,34^. While earlier clinical trials of DR5 agonistic antibodies showed minimal efficacy, they were generally well-tolerated ^17,18^. The poor clinical performance of these antibodies may be related to incomplete DR5 clustering or insufficient engagement of downstream signaling ^13,15–18^. Indeed, more potent multimeric antibodies, including INBRX-109, are now being evaluated in clinical trials. In contrast, DR5 CAR-T cells provide sustained engagement and simultaneous activation of multiple killing pathways, including granzyme/perforin release and FasL signaling, which may overcome the limitations of prior DR5-targeted approaches.

Importantly, we found no evidence of on-target off-tumor toxicity in either xenograft or syngeneic mouse models. This is encouraging, given that DR5 is detectable in some normal tissues. Several factors may help explain this safety profile. Normal cells often co-express decoy receptors such as DcR1 and DcR2, which can limit DR5-mediated apoptosis. Additionally, CAR- T cells may have limited access to or expansion in normal tissues, further reducing the risk of off-tumor effects. Ongoing clinical trials using DR5-targeted CAR-T cells in liver and breast cancers (NCT03941626, NCT06251544) have not yet reported severe toxicities, though published data remain limited.

Beyond direct cytotoxicity, DR5 CAR-T cells also altered the tumor immune landscape. MDSCs are well-known for suppressing T cell function and supporting tumor immune evasion ^35, 36 37^. In our study, DR5 agonistic CAR-T cells demonstrated the capacity to inhibit MDSCs, an important advantage in this challenging setting. In patient-derived melanoma organoids and slice cultures, which better recapitulate human tumor architecture and immunosuppression, DR5 CAR-T cells infiltrated tumor tissues and effectively depleted both M-MDSC and PMN-MDSC subsets ^28–31,38–40^, contributing to the increased numbers of Granzyme B+ and Ki-67+ CD8+ T cells in tumors, which indicates relief of immunosuppression and reactivation of tumor-resident immune cells. This immune activation is especially relevant given the known role of MDSCs in mediating resistance to checkpoint blockade and other therapies.

Despite these promising results, several questions remain. DR5 expression can vary across tumor types and even within tumors, which may impact treatment outcomes. Long-term persistence, safety, and antigen escape should be carefully evaluated in future studies. While our findings in preclinical models are encouraging, clinical validation will be critical to determine whether these benefits translate to patients.

In summary, DR5 CAR-T cells demonstrate a multifaceted mechanism of action, targeting both tumor cells and immunosuppressive myeloid populations. Their activity in immunocompetent models and human-derived tissues, along with a favorable safety profile, supports further development of DR5 CAR-T therapy as a candidate for solid tumor treatment.

## Methods

### Human DR5 CAR vector construction

We obtained four anti-human-DR5 antibody sequences from the International Immunogenetics Information System (IMGT, mAB-DB ID 183, 224, 234, and 348). The amino acid sequences of humanized DR5 scFvs were converted to cDNAs and custom synthesized by Integrated DNA Technologies, Inc. (CA, USA). The DNA fragments of the variable domain of heavy chain (VH) and light chain (VL) were fused and linked with a -(G4S)3- linker and flanked by a human-CD8 leader. These cDNAs were then cloned into an engineered pTRPE CAR encoding lentiviral backbone, which contains the EF1a promoter, CD8a leader, CD8a hinge extracellular domain, and transmembrane domain, followed by 4-1BB and CD3ζ endodomains, as previously described ^41,42^.

### Lentivirus packaging

CAR lentiviral vector packaging was carried out using HEK293T cells ^42^. A transfection mixture was prepared by combining 10 µg of the lentiviral vector plasmid, 7.5 µg of pMDLg/pRRE, 2.5 µg of pRSV-Rev, and 2.5 µg of pCMV-VSVG in 500 µL of serum-free RPMI medium. Premixed lipofectamine 2000 was added to the mixture and incubated for 20 minutes at room temperature. The transfection mix was added to the cells and incubated for 6-8 hours at 37°C in a 5% CO2 incubator. Half volume of full RPMI-1640 medium containing 10% FBS was added after transfection. After 24 and 48 hours, lentivirus-containing supernatant was collected, filtered, and concentrated overnight by centrifugation at 12,000 rpm. The final lentivirus particles were aliquoted and can be stored at -80°C until needed.

### Primary cells and cell lines

Human peripheral blood mononuclear cells (PBMCs) and monocytes from healthy donors were obtained from the Human Immunology Core at the Perelman School of Medicine at the University of Pennsylvania ^43^. Human melanoma cell lines A375, A2058, WM9, and 903 were obtained from Meenhard Herlyn’s laboratory (The Wistar Institute, Philadelphia, Pennsylvania, USA), and they were routinely tested for mycoplasma and DNA fingerprinted ^44^. Other cells were from ATCC.

### T cell expansion and CAR-T cell transduction

T-cell expansion was started from PBMCs using the method described ^42,45^. PBMCs were cultured with CD3/28 Danybeads at a 1:1 cell-to-bead ratio in RPMI1640 media supplemented with 10% FBS (HyClone; GE Healthcare, Utah, USA), 100 U/mL penicillin–streptomycin, 2 mM L-glutamine, 1/1000 2-mercaptoethanol (2-Me) (Gibco; Thermo Fisher Scientific, Massachusetts, USA) and 100 units/mL of recombinant human interleukin (IL)-2 (PeproTech, New Jersey, USA), new media added and supplemented every other day. The cells were incubated at 37°C with 5% CO2 for 24 hours before transduction with CAR lentiviral vectors at MOI=1. The medium and IL2 were replenished every 24-48 hours as required. After 7 days, the activated T cells were harvested for further purification, analysis, and downstream applications.

### MDSC culture

Isolated healthy donor PBMCs were used to culture MDSCs, with fresh cells used every time. The cells were plated at a density of 1-2 x 10^6 cells/mL in RPMI-1640 full medium and supplemented with 50 ng/ml of human GM-CSF and 20 ng/ml of human IL-6 in a T25 flask. Fresh culture medium was added every other day. After 7-10 days of culture, adherent cells were harvested and analyzed by flow cytometry to confirm MDSC phenotype, demonstrating that the cells were CD11b+CD33+ with other checkpoint markers ^46^. DR5 expression was assessed and compared to that of melanoma A375 cells using flow cytometry. This method was reliable and efficient for generating MDSCs for further experimentation.

### Melanoma xenografts

Animal procedures in this study were approved by the Institutional Animal Care and Use Committees at the University of Pennsylvania (804874) ^44^. All mice were bred and housed in a pathogen-free facility under controlled temperature (22 °C) and lighting (12:12 h light-dark cycle) and were fed a chow diet. Melanoma xenografts were established by injecting a single-cell suspension of A375 cells (2 million in 200 µL PBS) subcutaneously into the left flank of each nude mouse. Female nude mice from Jackson Laboratories were used in this study, and 5 mice were in each group. Treatment was initiated once tumors were palpable and had reached a size of approximately 50 mm^3^. Treatments were administered 5x10^6^ CAR+ T cells and same number of UTD cells in 100ul PBS as once or twice a week for 3 weeks. Tumor luminescence was calculated by measuring bioluminescence imaging ^44^. The instrument held 3 mice in one session. Tumor volume was measured using a caliber.

### Cytotoxicity assays

The Promega CytoTox 96 Non-Radioactive Cytotoxicity Assay kit (G1780; Promega, Wisconsin, USA) was utilized to measure the cytotoxicity of DR5 CAR T cells against melanoma cells and other cancer cells through lactate dehydrogenase (LDH) release, following previously described protocols ^43^. Target cells were seeded at a density of 4×10^4^ cells/well in 50 µL of standard growth medium in 96-well plates, while DR5 CAR-T cells were seeded at the indicated effector:target (E:T) ratio in the same volume simultaneously. After the stated time, the plates were briefly centrifuged, and 50 µL of supernatants were harvested for further analysis. The absorbance at 490 nm was measured using a BioTek Synergy HT reader (BioTek Instruments, Vermont, USA) following LDH activity detection. The percentage of cytotoxicity was calculated as (experimental−effector spontaneous−target spontaneous)/(target maximum−target spontaneous) ×100. Additionally, a luciferase-releasing assay system was employed to evaluate DR5 CAR-T cell cytotoxicity against target cells expressing luciferase. After incubation of target cells and DR5 CAR-T cells, the plates were briefly spun down, and the supernatant was carefully removed. Then, 25 µL of lysis buffer was added, and the plate was incubated on ice for 10 minutes before adding 100 µL of luciferin substrate of luciferase. The plate was read immediately using a luminometer. The percentage of cytotoxicity was calculated as (target maximum signal - experimental signal) / target maximum signal × 100.

### Flow cytometry

T cells and cancer cells were washed with a PBS solution containing 3% FBS. After washing, 0.5-1x10^6 cells were aliquoted into separate tubes and incubated with an antibody mixture in 100-150 ul of staining buffer for 20 minutes at 4°C in the dark. The cells were then washed once with the washing buffer and resuspended in 300-500 ul of washing buffer. If fixing was necessary, the cells were washed with PBS and fixed with a fixing buffer containing 1.5% PFA. Data acquisition was carried out using either BD LSR or Fortessa flow cytometers, and FlowJo software was used to analyze the data ^43^.

### Western blot

Western blotting was performed as described previously ^44,47^. Cells or extracellular vesicles (EVs) were lysed using RIPA buffer, and the protein concentrations were quantified using a BCA Protein Assay Kit (Bio-Rad, CA, USA). Subsequently, 20-50 μg of protein was separated by SDS-PAGE and e-transferred onto PVDF membranes. The membranes were then blocked with 5% non-fat milk at room temperature for 1 hour, followed by overnight incubation at 4 °C with primary antibodies. Afterward, the membranes were incubated with HRP-conjugated secondary antibodies (Cell Signaling Technology) at room temperature for 1 hour before being visualized using an ECL detection reagent.

### Reverse phase protein array assays (RPPA)

RPPA sample preparation and protocol, as previously described ^43,44^. Briefly, protein lysates were prepared using a lysis buffer and 4× SDS sample buffer, with added protease and phosphatase inhibitors. The lysates were extracted from frozen tumors and analyzed using the RPPA platform at the MD Anderson Functional Proteomics Core facility. Detailed information on the RPPA method, data normalization, and the antibodies used is available on the core facility’s website. Heatmaps were generated using Cluster and then analyzed using R and The Database for Annotation, Visualization, and Integrated Discovery (DAVID) ^48,49^.

### Computational analysis of scFv binding affinity to DR5

To model the single-chain variable fragment (scFv), the variable light and heavy chains of each antibody were obtained from the International Immunogenetics Information System ^50,51^. These, along with the linker sequence, were then entered into the Antibody Structure Prediction Panel of Maestro suite (2023-2) by Schrödinger [Schrödinger, L., and Warren DeLano. "PyMOL."

(2023)]. The Antibody Structure Prediction Panel employed a homology search for the framework region to model the scFvs, utilizing the Enhanced Clothia numbering scheme. The highest-scoring homolog framework, defined by the average of the heavy and light chain similarity scores of the framework region to that of the inputted sequence, was chosen for each scFv. After the framework region selection, the scFv models were submitted for the complementarity-determining region (CDR) loop modeling, where antibody databases were searched for loops of the same length as those in the query sequence. The loops are then clustered structurally. The modeled scFv is inspected in the workspace where the hydrogen bonding network was optimized within the structure, followed by a minimization step and water removal.

Once homology modeling was done, the scFvs were further prepared for docking studies using PDB-Tools (v2.3.1) webserver ^52^. The linker was extracted, leaving only the variable domain of the scFvs, followed by sequential renumbering of the residue sequence. A final clean-up was conducted to remove any irrelevant lines that could disturb the residue renumbering. Once the .pdb file for each scFv was prepared, the high-resolution 2.2 Å crystal structure of DR5 (PDB ID:1D4V) ^53^ was obtained from the online RCSB Protein Data Bank as .pdb files. It was then opened on PyMOL (2.5.5), where water molecules and any non-applicable bound ligands were removed. The final structures were then exported as a .pdb file to be used for molecular docking studies on the web server, High Ambiguity Driven protein-protein Docking (HADDOCK) (v2.4) ^54,55^. A loose definition of the epitope and paratope residues were specified to guide the docking.

### Expression of mouse-anti-human DR5 recombinant antibodies

To generate mouse-anti-human DR5 recombinant antibodies, the DR5 CAR scFv cDNAs were cloned into pFUSEss-CHIg-mG2a and pFUSE2ss-CLIg-mK vectors from Invivogene, while the anti-hDR5 VL or VH cDNAs were cloned into pFUSE2ss-CLIg-mK or pFUSEss-CHIg-mG2a vectors, respectively ^42^. FreeStyle 293-F cells were cultured in FreeStyle 293 Expression Medium under shaking conditions at 90 rpm and in an incubator at 37°C and 5% CO2 until they reached a density of 2-3 x 10^6 cells/mL. The paired DR5 pFUSEss-CHIg/CLIg plasmids were transfected into the cells using 293fectin Transfection Reagent. After 24-48 hours of growth, the supernatant was collected, and cell debris was removed by centrifugation at 10,000 × g for 20 minutes. Recombinant antibody concentrations were determined using a BCA Protein Assay Kit and SDS-PAGE/Silver Stain, and the purified antibodies were stored at -20°C or 4°C until further use.

### On-Cell-Western (OCW) assay

OCW assays were performed at the Wistar Institute. A375 melanoma cells with overexpressed GFP and DR5 were utilized as target cells for the OCW assay. For the live cell assay, 8,000 cells were seeded and allowed to attach for 24 hours. Antibodies (Abs) were dissolved in culture medium at dilutions of 1:2, 1:4, and 1:8. A total of 30 µL of the Ab solution was added to each well and incubated at 37°C for 30 minutes. The cell media was removed, and 50 µL of 4% paraformaldehyde was added to fix the cells. The plate was incubated at room temperature for 20 minutes without agitation. After washing the plate twice with TBS, 50 µL of Intercept Blocking Buffer was added to each well and incubated for 1 hour at room temperature with moderate shaking to allow for blocking. A fluorescently labeled secondary antibody, IRDye 800 CW, was diluted in 0.2% Tween-20 Intercept Blocking Buffer at a ratio of 1:800, and LI-COR CellTag 700 Stain was diluted at 1:500. 15 µL of the diluted secondary antibody solution with CellTag 700 Stain was added to testing wells, while 15 µL of the secondary antibody solution without CellTag 700 Stain was added to background wells. The plate was incubated on a plate shaker for 45 minutes, protecting it from light. The plate was washed 5 times with 1X TBS with 0.1% Tween- 20 washing solution, and the final wash solution was removed by tapping or blotting the plate. The plate was then scanned on the LI-COR Odyssey CLx with the 700 and 800 nm channels using an Odyssey® Imager. The data was analyzed with Image Studio under the On-Cell Western Software protocol and graphed using Prism GraphPad 8. The experiments were performed in triplicate.

### Extracellular vesicle preparation and characterization

To isolate exosomes from cell culture, the cell culture supernatant is collected and centrifuged at 300 × g for 10 minutes to remove any cells or debris. The resulting supernatant is then transferred to ultracentrifuge tubes and centrifuged at 2,000 × g for 10 minutes to remove any remaining cell debris. After filtering the supernatant through a 0.22 µm filter, it is centrifuged at 100,000 × g for 2 hours to pellet the exosomes. The supernatant is then discarded, and the exosome pellet is washed and resuspended in a small volume of PBS or another appropriate buffer, followed by another round of ultracentrifugation at 100,000 × g for 2 hours. Finally, the supernatant is discarded, and the exosome pellet is resuspended in a small volume of PBS or another appropriate buffer for downstream applications. The NanoSight NS300 (Malvern Instruments), which comes with particle-tracking software and fast video capture capabilities, was used to measure the size and concentration of sEVs obtained from Sup-T cell culture supernatants ^33,47^.

### Bicellular spheroid culturing

The method for generating bicellular spheroids and co-culture was previously described ^38,39,43,56^. In summary, 50 μl of 1.5% agarose gel (A9539, Sigma-Aldrich, St. Louis, Missouri) was used to prime the culturing well of a flat bottom 96-well tissue culture multiplate to form an unattached support matrix. Bicellular spheroids were created by seeding 3 x 10^3^ A375 melanoma cells and 6 x 10^3^ BJ fibroblasts (tumor cells:fibroblasts = 1:2) or 3 x 10^3^ A375 melanoma cells and the same number of MDSCs (tumor cells:MDSCs = 1:1) into the agarose-primed wells. After 48 hours of co-culturing, the spheroids were checked under a microscope and then co-cultured with 3 x 10^3^ CAR-T cells (tumor cells:effectors = 1:1). Prior to fluorescence microscope scanning after 24 hours of co-culturing, Hoechst33342 (1μg/ml) and PI (1μg/ml) were stained. FUJI imaging was used to analyze imaging data and all experiments were set up in triplicate.

### Murine DR5 CAR construct and murine CAR-T cell manufacture

The murine DR5 CAR is designed based on scFv sequence of the DR5 M5-1 monoclonal antibody ^26^ and the DNA construct has traceable BFP and nLuc as we previously described ^27^. Mouse CAR T cells were generated using a retroviral transduction protocol. First, Plat-E cells were transfected with a pMMLV-based vector encoding the CAR construct along with fluorescent and bioluminescent reporters. The resulting retroviral supernatant was harvested and used to transduce mouse CD3+ T cells isolated from splenocytes and activated with plate-bound anti- CD3/CD28 antibodies. RetroNectin-coated plates and centrifugation were used to facilitate viral transduction. After transduction, CAR T cells were expanded in IL-2-containing media for up to six days. Transduction efficiency was confirmed by flow cytometry using both reporter fluorescence and CAR-specific staining.

### Melanoma patient-derived organoids (MPDOs)

Fresh melanoma tissues were obtained from the University of Pennsylvania Hospital Department of Pathology and Laboratory Medicine. All procedures were approved by the Institutional Review Board at the University of Pennsylvania. The fresh tissues were maintained in RPMI-1640 medium until processing. The tissue specimen blocks and organoids used in this study were prepared as previously described ^39,40^. To optimize the growth of melanoma organoids, we developed a FBS-free medium specifically designed for melanoma cells independently, which comprised of 50% M594 media, 50% Advanced DMEM media, and 1% each of B27, NEAA, and P/S.

### Patient-derived melanoma slice culture (MSC)

The fresh tumor tissues were kept on ice and sliced using a vibrating microtome. Tissue preparation was performed according to a previously described protocol ^28^. Briefly, the tissues were collected and embedded in 2% agarose for slicing. After the agarose solidified, it was fixed to a specimen disc and sliced using Compresstome VF-310-0Z at a thickness of 300 μm. The tissue slices were placed into a Millicell insert (PIHP01250, Millipore, MO, USA) in a 24-well plate and cultured in RPMI 1640 medium with 10% FBS, 2 mM L-glutamine (A2916801, Gibco), 10 mM HEPES (15630080, Gibco), 1 mM sodium pyruvate (11360070, Gibco), 5.5 µM 2- mercaptoethanol, 1% penicillin–streptomycin, and 200 IU/mL IL-2.

### Immunofluorescence and live imaging

Immunofluorescence and live image pictures were captured and analyzed using a Zeiss microscope with 10x and 20x objective lenses. To visualize the nuclei and cell viability, cells were stained with Hoechst33342 and PI, respectively. Depending on the specific experiments, T-cells were stained with CFSE or CellTrace Far Red. Zeiss ZEN software was employed to analyze the fluorescent images, which were synthesized into video files. Representative images were used to present the data, while statistical analyses were based on at least three independent datasets.

### Statistical analysis

Statistical analysis was performed using GraphPad (Prism software package version) and Microsoft Excel software. Data are presented as mean±SEM, and significant differences were examined with Student’s t-test. A P value of <0.05 was considered statistically significant.

## Supporting information

Supplemental Movie 1.Co-Culture MPDO with UTD and DR5 CAR-T Cells

## Acknowledgments

The authors thank the Penn Cytomics and Cell Sorting Shared Resource Laboratory at the Perelman School of Medicine at the University of Pennsylvania, especially Shifu Tian, for providing protocols for CAR-T cell purification. We thank the Functional Proteomics RPPA Core at MD Anderson Cancer Center for performing the RPPA assay and raw data cleaning and analysis. We also thank Lily Lu at The Wistar Institute Molecular Screening and Protein Expression Core Facility for testing the affinity of DR5 antibodies.

## Author contributions

XX and HW conceived the project and designed the study. XX, HW GK, RA, LS, and SL collected and analyzed the samples. HW, SL, PS, XZ, QZ, HM, JY, YG, LH, and FL performed the experiments. HW, SL and PS manufactured DR5 CAR T cells. FL performed the bioinformation and analysis. BG performed the antibody computational analysis. YG designed the graphical abstract. HW, JS, FL, FM, CZ, MM, YF, and CJ contribute to CAR-T cell development. XX, AH, TM, YF, CJ, DG, WG, and MH provided insights into target and CAR-T cell development. HW, SL, and XX wrote the manuscript.

## Data and materials availability

All data needed to evaluate the conclusions of the paper are present in the paper and/or the Supplementary Materials. DNA constructs can be provided by The Trustees of the University of Pennsylvania pending scientific review and a completed material transfer agreement.

## Supplementary Figures

**Fig. S1.**
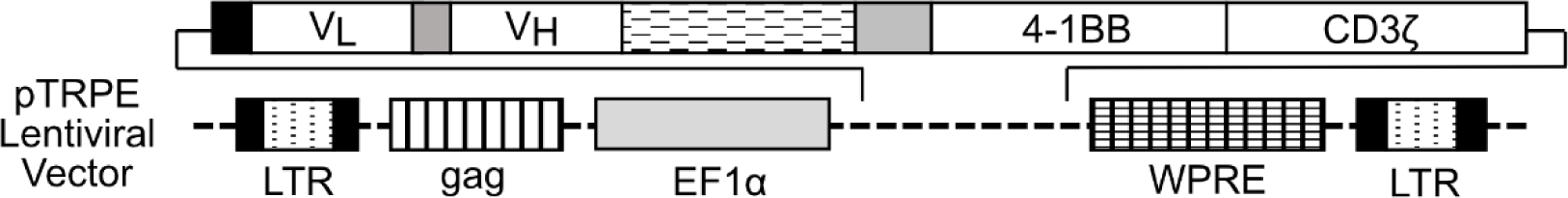
**Schematic of pTRPE lentiviral vector for lentiviral DR5 CAR expression.**

**Fig. S2.**
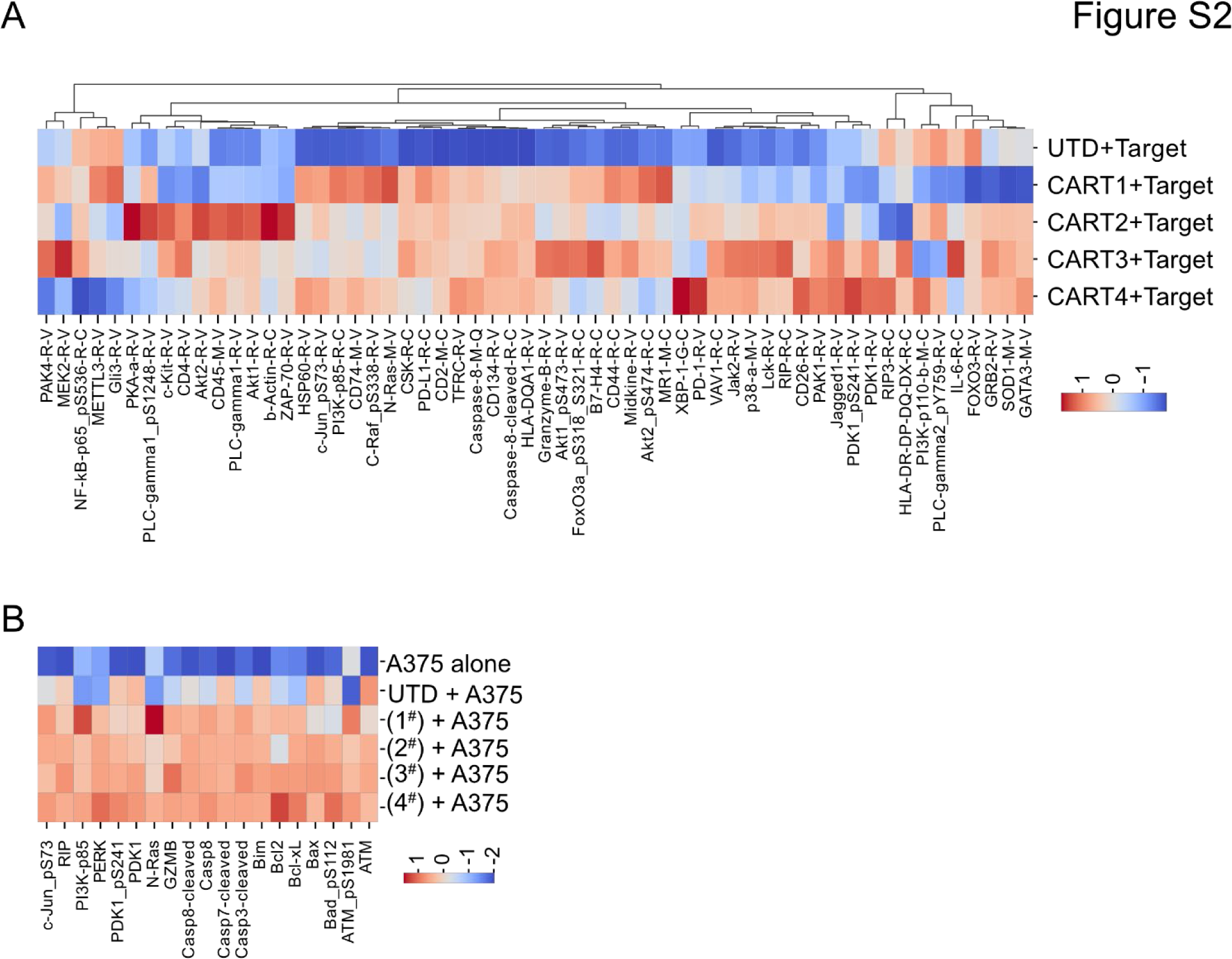
DR5 CAR-T cell activation after incubation with target cells. (A) Heat map of T cell activation gene expression after incubation of DR5 CAR-T cells with A375 melanoma cells using RPPA. Data shown as average values of 5 samples (n=5). (B) Heat map of cell death and apoptosis-associated protein expression using RPPA after co-culture of A375 melanoma cells with CAR-T cells for 4 hours, n=5.

**Fig. S3.**
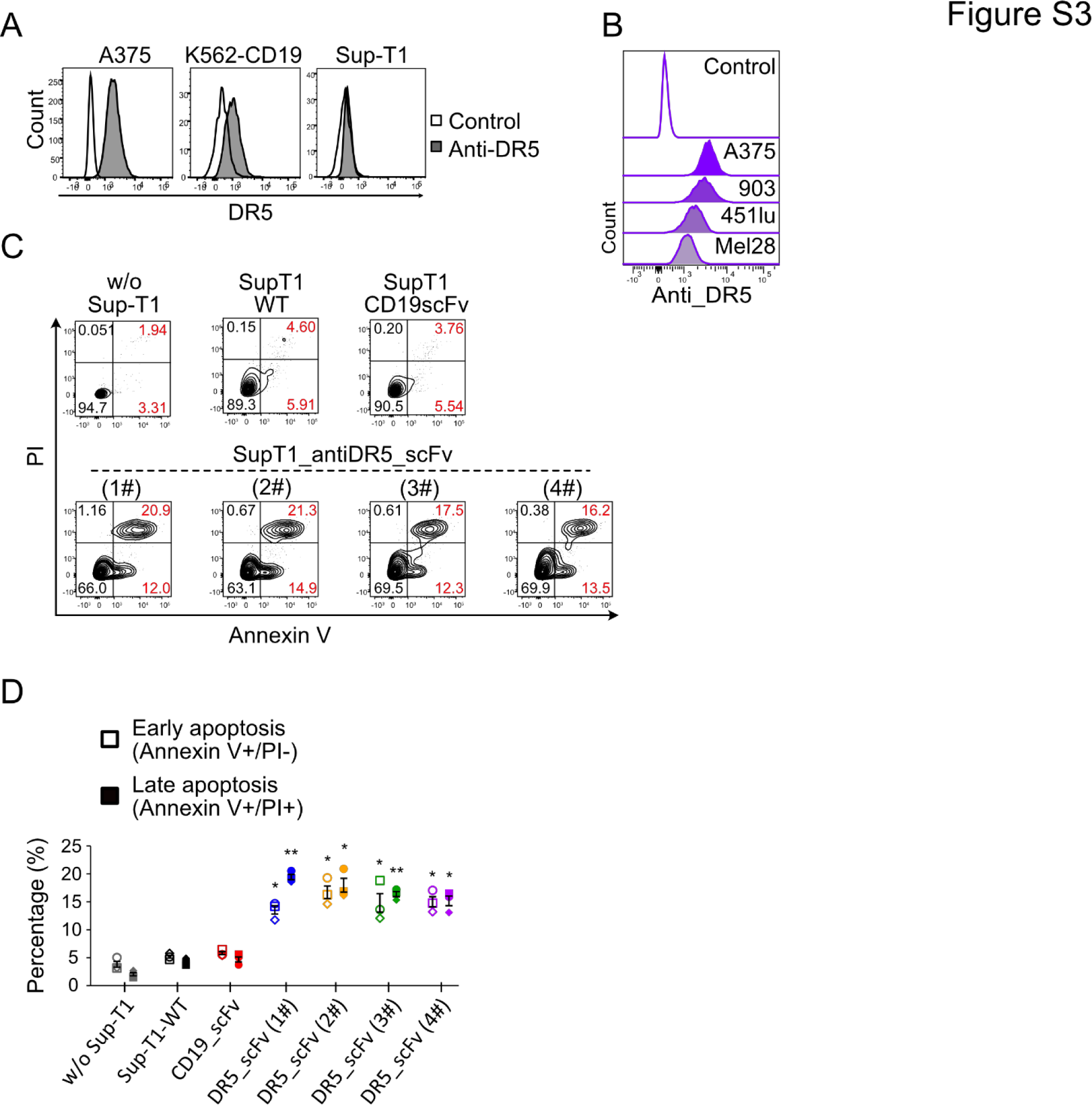
Effects of DR5-CAR-SupT1 cells on K562-CD19 cells. (A) DR5 expression levels in different cell lines. DR5 expression on A375, K562-CD19, and SupT1 cells was measured by FACS. (B) DR5 expression in a panel of melanoma cell lines by FACS analysis. (C-D) Effects of DR5-CAR-SupT1 cells on K562-CD19 cells. SupT1 or DR5 CAR-SupT1 cells are cocultured with K562-CD19 cells at an E:T ratio of 1:1 for 16 hrs. PI and Annexin V were used to measure cell apoptosis. Representative flow chart after coculture of SupT1 cells with K562-CD19 (C). Statistical analysis of apoptosis and cell death of K562-CD19 cells after treatment, n=3 (D). All data are shown as Mean±S.E.M., *P<0.05, **P<0.01, ***P<0.001, n.s., no statistically significant difference.

**Fig. S4.**
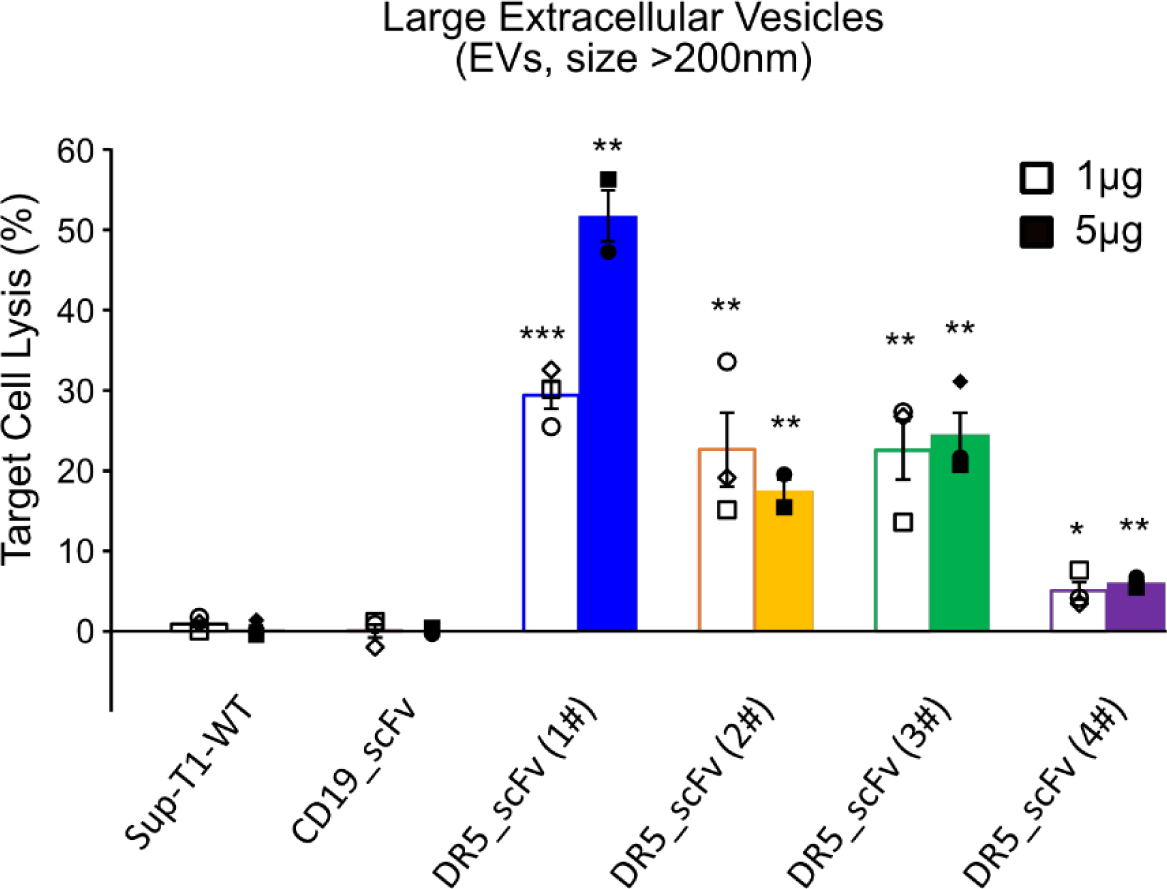
Cytotoxicity of large EVs derived from DR5 CAR-SupT1 cells to A375 cells. Luciferase-based assays were performed after coculture of large EVs (1 μg/ml or 5 μg/ml) from DR5 CAR-SupT1 with A375 cells for 16 hrs, n=3. All data are shown as Mean±S.E.M., *P<0.05, **P<0.01, ***P<0.001, n.s., no statistically significant difference.

**Fig. S6.**
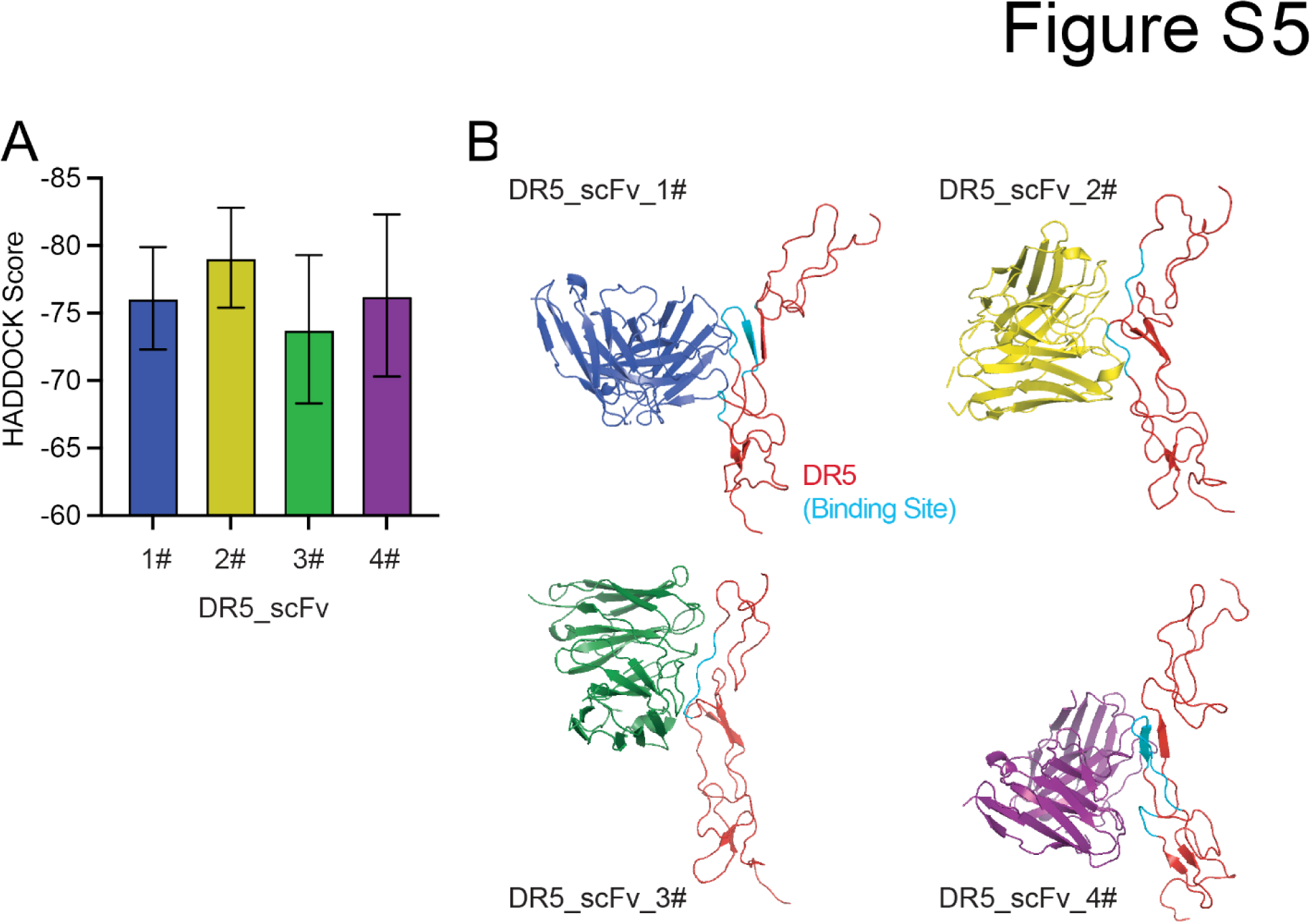
Computational analysis of DR5 scFvs binding to their targets. (A) HADDOCK scores of affinities of DR5 scFvs to their targets. (B) Computational models of DR5 scFvs binding to DR5 protein. The binding sites are highlighted with a teal color.

**Fig. S6.**
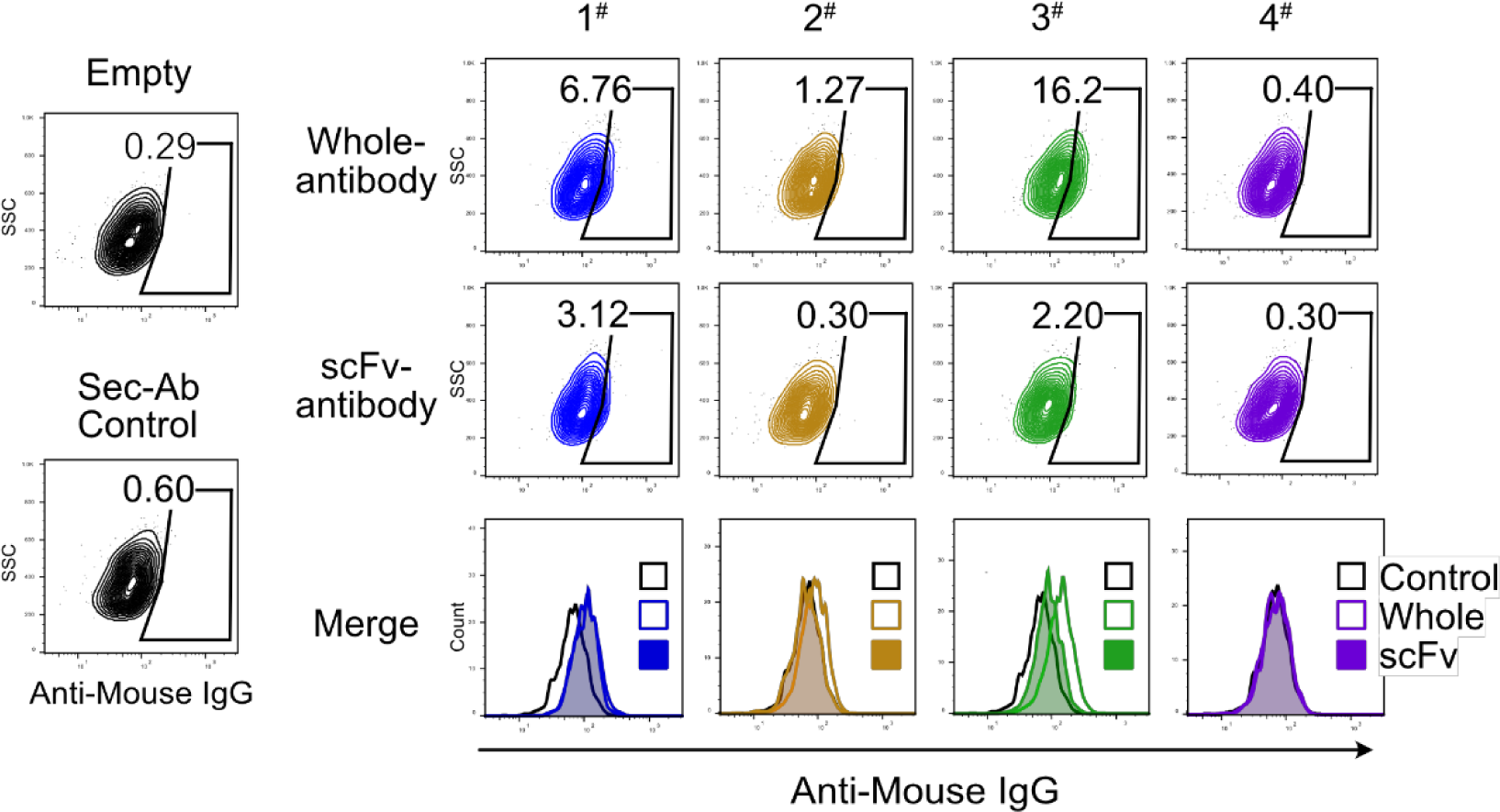
Representative flow cytometry data showing the DR5 antibody binding to A375 cells. Four different DR5 whole and scFv antibodies (4μg) were used in the assays.

**Fig. S7.**
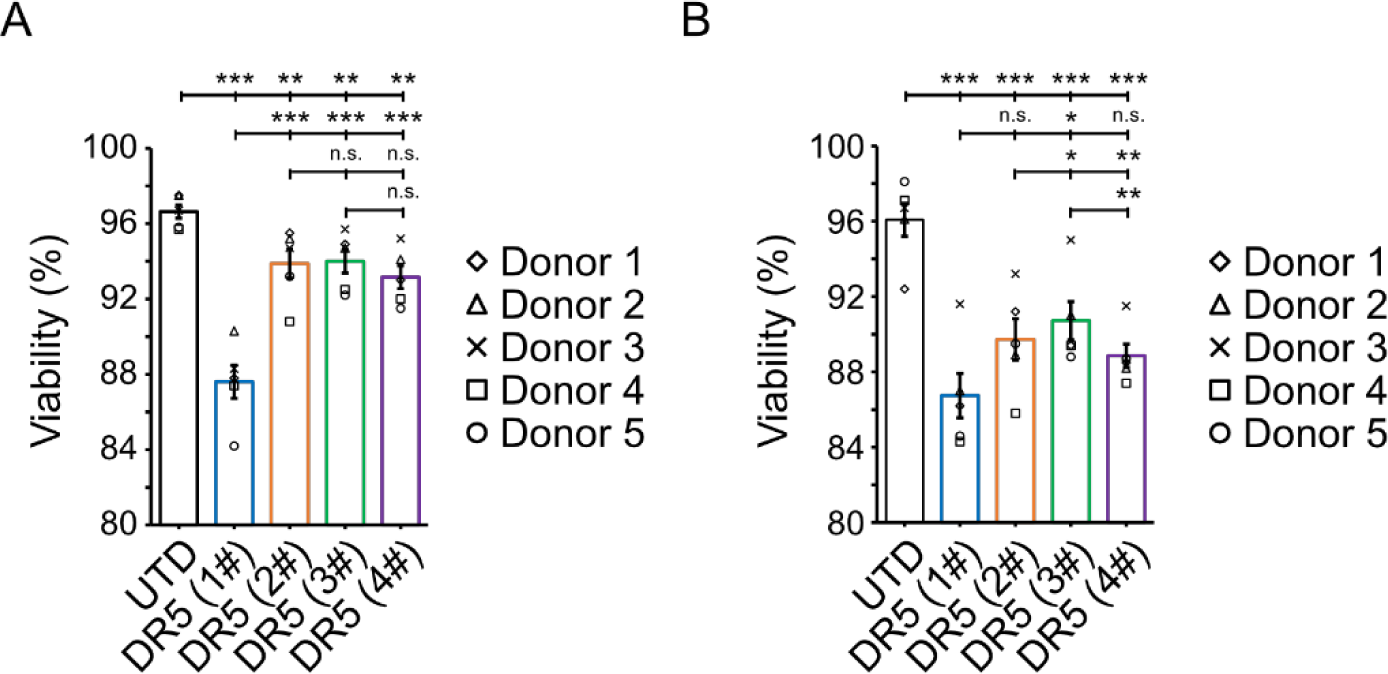
CAR-T cell viability after expansion and activation. (A) Viability of cultured DR5 CAR-T cells after expansion for 4 days, n=5. (B) Viability of cultured DR5 CAR-T cells after expansion for 3 days and cocultured with target cells (A375) for 24 hours, n=5. All data are shown as Mean±S.E.M., *P<0.05, **P<0.01, ***P<0.001, n.s., no statistically significant difference.

**Fig. S8.**
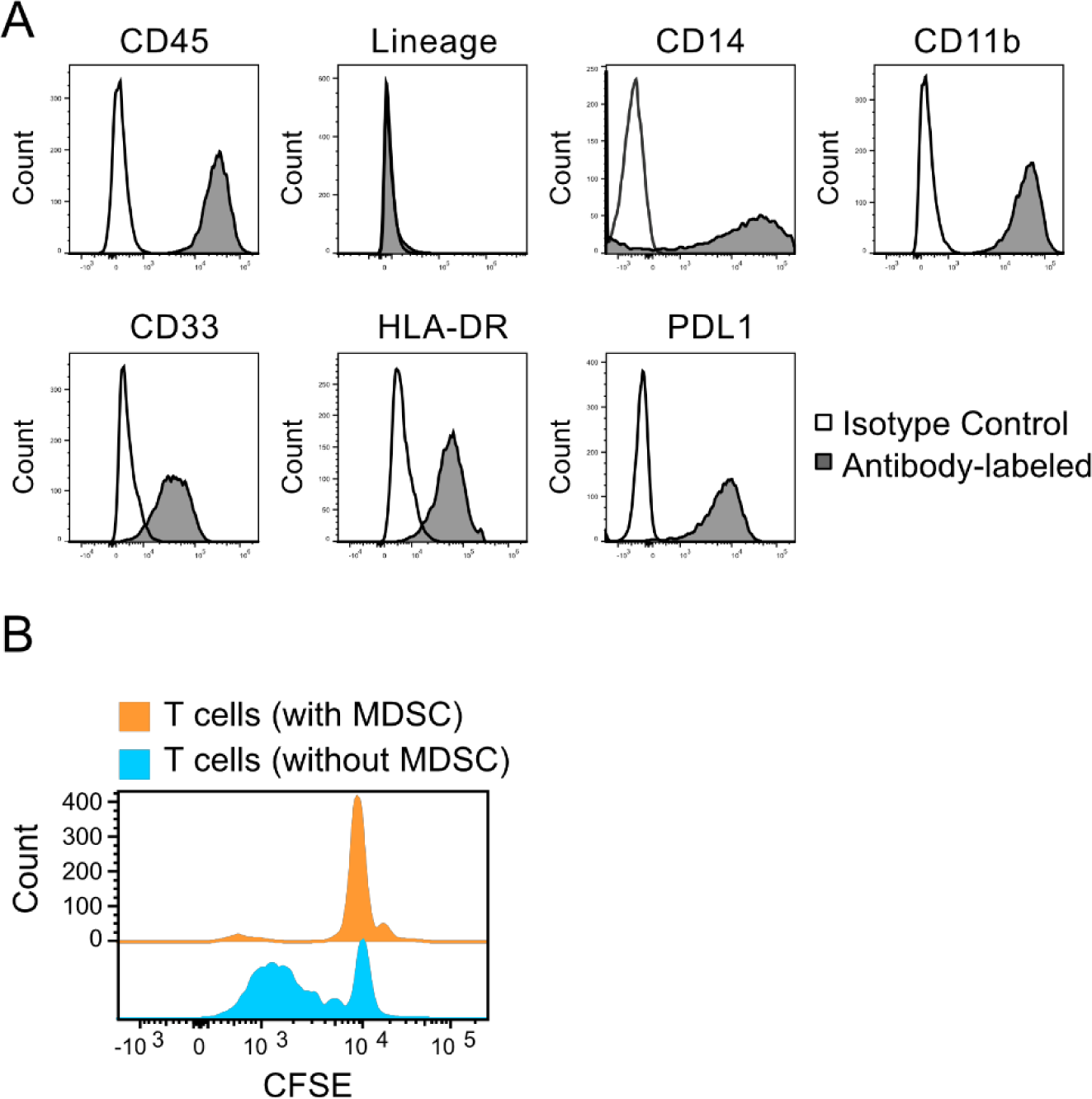
MDSCs inhibit T cell proliferation. (A) PBMC-monocyte-derived MDSCs were typically identified as CD11b^+^CD33^+^PDL1^hi^ cells by FACS. They also express HLD-DR and CD14. Representative flow charts of induced MDSCs. (B) Induced MDSCs inhibited T-cell proliferation. CFSE staining data after co-culturing T cells and induced MDSCs for 3 days.

**Fig. S9.**
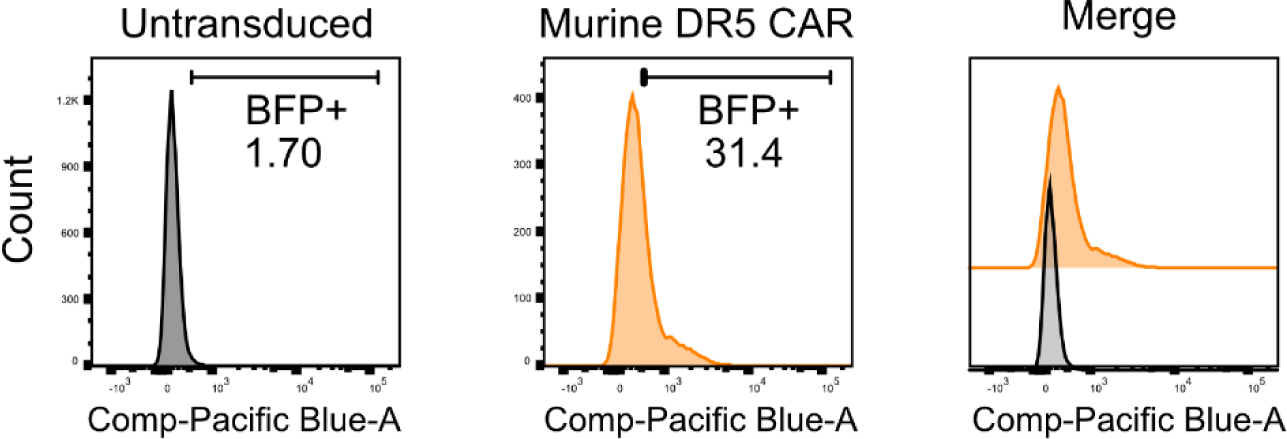
Representative flow charts showing murine DR5 CAR-T cells express BFP.

**Fig. S10.**
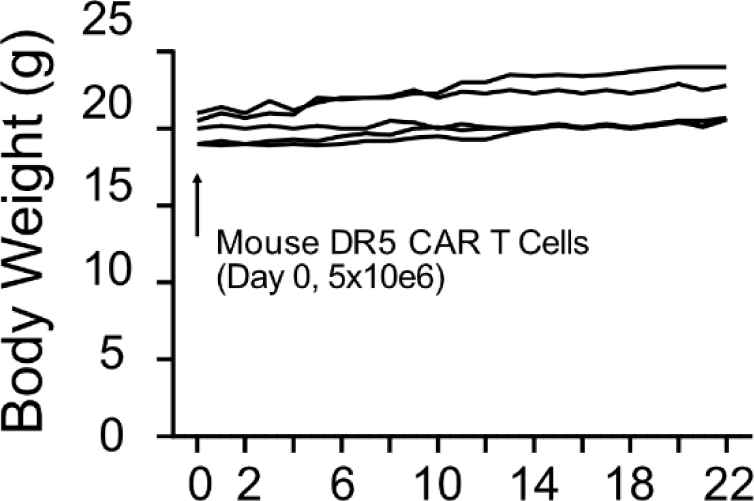
Body weight of C57 mice injected with murine DR5 CAR-T cells. Non-tumor bearing mice were injected with 5x10^6^ murine DR5 CAR-T cells via the tail veins. Body weight of the treated mice was measured twice a week, n=5.

**Fig. S11.**
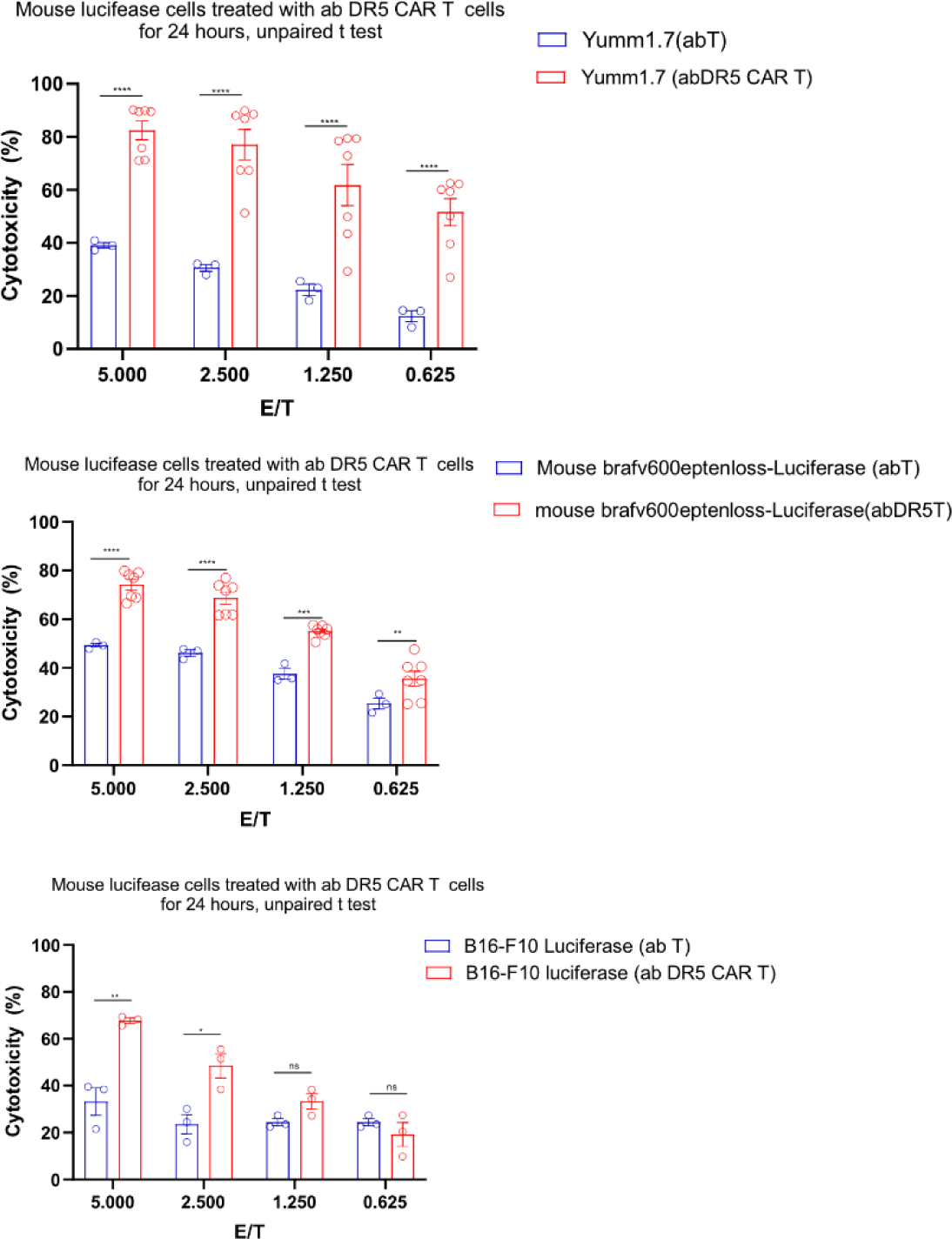
Murine DR5 CAR-T cells are cytotoxic to murine melanoma cells. Luciferase cytotoxicity assay after coculture of murine DR5 CAR-T cells with cancer cells for 24 hours at indicated E:T ratio, n=3. All data are shown as mean±SEM. *P<0.05, **P<0.01, ***P<0.001, n.s., no statistically significant difference.

**Fig. S12.**
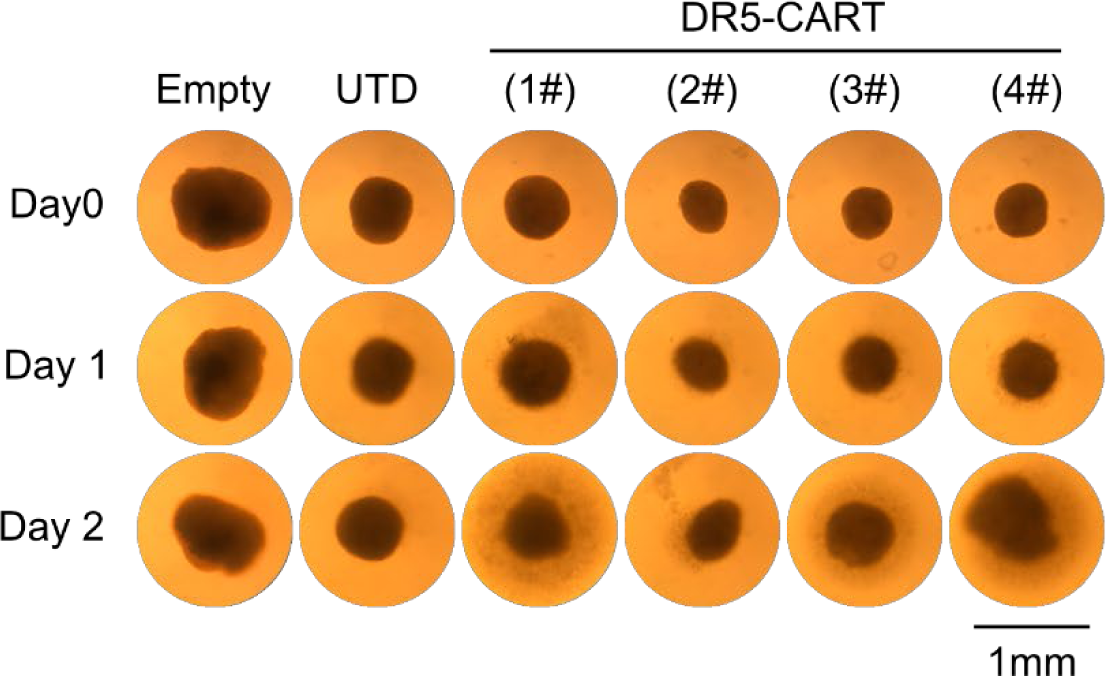
DR5 CAR-T cells induced tumor lysis in the MPDOs. Representative bright field images of MPDOs from four melanoma patients cocultured with DR5 CAR-T cells for 48 hours showing organoid disintegration.

**Fig. S13.**
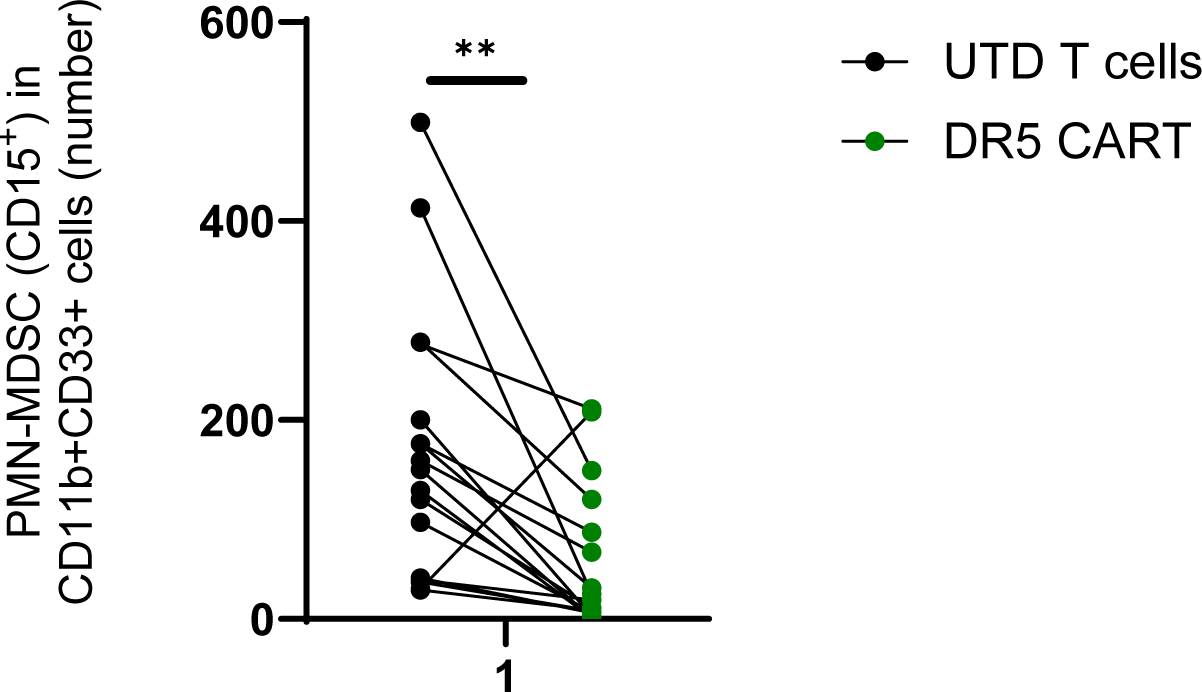
DR5 CAR-T cells inhibit PMN-MDSCs in the MPDOs. MPDOs from four melanoma patients are incubated with labeled UTD or CAR-T cells for 48 hours. After treatment, the MPDOs were collected and dissociated into single cells for FACS analysis. CD15+CD1b+CD33+ PMN-MDSCs were significantly decreased after CAR-T cell treatment. n=17, and each dot represents an independent well.

## References

1. Chen, T., Wang, M., Chen, Y., and Liu, Y. (2024). Current challenges and therapeutic advances of CAR-T cell therapy for solid tumors. Cancer Cell International 24, 133.

2. Marofi, F., Motavalli, R., Safonov, V.A., Thangavelu, L., Yumashev, A.V., Alexander, M., Shomali, N., Chartrand, M.S., Pathak, Y., and Jarahian, M. (2021). CAR T cells in solid tumors: challenges and opportunities. Stem cell research & therapy 12, 1–16.

3. Martinez, M., and Moon, E.K. (2019). CAR T Cells for Solid Tumors: New Strategies for Finding, Infiltrating, and Surviving in the Tumor Microenvironment. Front Immunol 10, 128. 10.3389/fimmu.2019.00128.

4. Albelda, S.M. (2024). CAR T cell therapy for patients with solid tumours: key lessons to learn and unlearn. Nature Reviews Clinical Oncology 21, 47–66.

5. Singleton, D.C., Macann, A., and Wilson, W.R. (2021). Therapeutic targeting of the hypoxic tumour microenvironment. Nature reviews Clinical oncology 18, 751–772.

6. Mao, X., Xu, J., Wang, W., Liang, C., Hua, J., Liu, J., Zhang, B., Meng, Q., Yu, X., and Shi, S. (2021). Crosstalk between cancer-associated fibroblasts and immune cells in the tumor microenvironment: new findings and future perspectives. Molecular cancer 20, 1–30.

7. Haist, M., Stege, H., Grabbe, S., and Bros, M. (2021). The functional crosstalk between myeloid-derived suppressor cells and regulatory T cells within the immunosuppressive tumor microenvironment. Cancers 13, 210.

8. Russo, M., and Nastasi, C. (2022). Targeting the tumor microenvironment: a close up of tumor-associated macrophages and neutrophils. Frontiers in Oncology 12, 871513.

9. Liu, G., Rui, W., Zhao, X., and Lin, X. (2021). Enhancing CAR-T cell efficacy in solid tumors by targeting the tumor microenvironment. Cellular & Molecular Immunology 18, 1085–1095.

10. Hou, A.J., Chen, L.C., and Chen, Y.Y. (2021). Navigating CAR-T cells through the solid- tumour microenvironment. Nature reviews Drug discovery 20, 531–550.

11. Condamine, T., Kumar, V., Ramachandran, I.R., Youn, J.I., Celis, E., Finnberg, N., El- Deiry, W.S., Winograd, R., Vonderheide, R.H., English, N.R., Knight, S.C., et al. (2014). ER stress regulates myeloid-derived suppressor cell fate through TRAIL-R-mediated apoptosis. J Clin Invest 124, 2626–2639. 10.1172/JCI74056.

12. Carneiro, B.A., and El-Deiry, W.S. (2020). Targeting apoptosis in cancer therapy. Nature reviews Clinical oncology 17, 395–417.

13. Yuan, X., Gajan, A., Chu, Q., Xiong, H., Wu, K., and Wu, G.S. (2018). Developing TRAIL/TRAIL death receptor-based cancer therapies. Cancer and Metastasis Reviews 37, 733–748.

14. Daniels, R.A., Turley, H., Kimberley, F.C., Liu, X.S., Mongkolsapaya, J., Ch’En, P., Xu, X.N., Jin, B., Pezzella, F., and Screaton, G.R. (2005). Expression of TRAIL and TRAIL receptors in normal and malignant tissues. Cell research 15, 430–438.

15. Pimentel, J.M., Zhou, J.-Y., and Wu, G.S. (2023). The role of TRAIL in apoptosis and immunosurveillance in cancer. Cancers 15, 2752.

16. Forero, A., Bendell, J.C., Kumar, P., Janisch, L., Rosen, M., Wang, Q., Copigneaux, C., Desai, M., Senaldi, G., and Maitland, M.L. (2017). First-in-human study of the antibody DR5 agonist DS-8273a in patients with advanced solid tumors. Investigational New Drugs 35, 298–306.

17. Snajdauf, M., Havlova, K., Vachtenheim Jr, J., Ozaniak, A., Lischke, R., Bartunkova, J., Smrz, D., and Strizova, Z. (2021). The TRAIL in the treatment of human cancer: an update on clinical trials. Frontiers in molecular biosciences 8, 628332.

18. Dominguez, G.A., Condamine, T., Mony, S., Hashimoto, A., Wang, F., Liu, Q., Forero, A., Bendell, J., Witt, R., and Hockstein, N. (2017). Selective targeting of myeloid-derived suppressor cells in cancer patients using DS-8273a, an agonistic TRAIL-R2 antibody. Clinical Cancer Research 23, 2942–2950.

19. Gabrilovich, D.I. (2017). Myeloid-derived suppressor cells. Cancer immunology research 5, 3–8.

20. Weber, R., Fleming, V., Hu, X., Nagibin, V., Groth, C., Altevogt, P., Utikal, J., and Umansky, V. (2018). Myeloid-derived suppressor cells hinder the anti-cancer activity of immune checkpoint inhibitors. Frontiers in immunology 9, 1310.

21. Herrmann, I.K., Wood, M.J.A., and Fuhrmann, G. (2021). Extracellular vesicles as a next-generation drug delivery platform. Nature nanotechnology 16, 748–759.

22. Nederveen, J.P., Warnier, G., Di Carlo, A., Nilsson, M.I., and Tarnopolsky, M.A. (2021). Extracellular vesicles and exosomes: insights from exercise science. Frontiers in physiology 11, 604274.

23. Flugel, C.L., Majzner, R.G., Krenciute, G., Dotti, G., Riddell, S.R., Wagner, D.L., and Abou-el-Enein, M. (2023). Overcoming on-target, off-tumour toxicity of CAR T cell therapy for solid tumours. Nature Reviews Clinical Oncology 20, 49–62.

24. Olson, M.L., Mause, E.R.V., Radhakrishnan, S.V., Brody, J.D., Rapoport, A.P., Welm, A.L., Atanackovic, D., and Luetkens, T. (2022). Low-affinity CAR T cells exhibit reduced trogocytosis, preventing rapid antigen loss, and increasing CAR T cell expansion. Leukemia 36, 1943–1946.

25. Duan, Y., Chen, R., Huang, Y., Meng, X., Chen, J., Liao, C., Tang, Y., Zhou, C., Gao, X., and Sun, J. (2022). Tuning the ignition of CAR: optimizing the affinity of scFv to improve CAR-T therapy. Cellular and Molecular Life Sciences 79, 14.

26. Takeda, K., Yamaguchi, N., Akiba, H., Kojima, Y., Hayakawa, Y., Tanner, J.E., Sayers, T.J., Seki, N., Okumura, K., and Yagita, H. (2004). Induction of tumor-specific T cell immunity by anti-DR5 antibody therapy. Journal of Experimental Medicine 199, 437–448.

27. Zhang, D., Krimitza, E., Han, K., Su, R., Xu, D.J., Xu, J.R., Gong, Y., and Fan, Y. (2024). Protocol to generate traceable CAR T cells for syngeneic mouse cancer models. STAR protocols 5, 102898.

28. Kenerson, H.L., Sullivan, K.M., Labadie, K.P., Pillarisetty, V.G., and Yeung, R.S. (2021). Protocol for tissue slice cultures from human solid tumors to study therapeutic response. STAR protocols 2, 100574.

29. Dimou, P., Trivedi, S., Liousia, M., D’Souza, R.R., and Klampatsa, A. (2022). Precision-cut tumor slices (PCTS) as an ex vivo model in immunotherapy research. Antibodies 11, 26.

30. Kenerson, H.L., Sullivan, K.M., Seo, Y.D., Stadeli, K.M., Ussakli, C., Yan, X., Lausted, C., Pillarisetty, V.G., Park, J.O., and Riehle, K.J. (2020). Tumor slice culture as a biologic surrogate of human cancer. Annals of translational medicine 8.

31. Sivakumar, R., Chan, M., Shin, J.S., Nishida-Aoki, N., Kenerson, H.L., Elemento, O., Beltran, H., Yeung, R., and Gujral, T.S. (2019). Organotypic tumor slice cultures provide a versatile platform for immuno-oncology and drug discovery. Oncoimmunology 8, e1670019.

32. Voabil, P., de Bruijn, M., Roelofsen, L.M., Hendriks, S.H., Brokamp, S., van den Braber, M., Broeks, A., Sanders, J., Herzig, P., and Zippelius, A. (2021). An ex vivo tumor fragment platform to dissect response to PD-1 blockade in cancer. Nature medicine 27, 1250–1261.

33. Guo, Y., Wang, H., Liu, S., Zhang, X., Zhu, X., Huang, L., Zhong, W., Guan, L., Chen, Y., and Xiao, M. (2025). Engineered extracellular vesicles with DR5 agonistic scFvs simultaneously target tumor and immunosuppressive stromal cells. Science Advances 11, eadp9009.

34. Buchsbaum, D.J., Zhou, T., and LoBuglio, A.F. (2006). TRAIL receptor-targeted therapy. Future Oncology 2, 493–508.

35. Grover, A., Sanseviero, E., Timosenko, E., and Gabrilovich, D.I. (2021). Myeloid-derived suppressor cells: a propitious road to clinic. Cancer discovery 11, 2693–2706.

36. Hegde, S., Leader, A.M., and Merad, M. (2021). MDSC: Markers, development, states, and unaddressed complexity. Immunity 54, 875–884.

37. Li, T., Liu, T., Zhu, W., Xie, S., Zhao, Z., Feng, B., Guo, H., and Yang, R. (2021). Targeting MDSC for immune-checkpoint blockade in cancer immunotherapy: current progress and new prospects. Clinical Medicine Insights: Oncology 15, 11795549211035540.

38. Ou, L., Wang, H., Huang, H., Zhou, Z., Lin, Q., Guo, Y., Mitchell, T., Huang, A.C., Karakousis, G., and Schuchter, L. (2022). Preclinical platforms to study therapeutic efficacy of human γδ T cells. Clinical and Translational Medicine 12, e814.

39. Ou, L., Liu, S., Wang, H., Guo, Y., Guan, L., Shen, L., Luo, R., Elder, D.E., Huang, A.C., and Karakousis, G. (2023). Patient-derived melanoma organoid models facilitate the assessment of immunotherapies. EBioMedicine 92.

40. Jacob, F., Salinas, R.D., Zhang, D.Y., Nguyen, P.T., Schnoll, J.G., Wong, S.Z.H., Thokala, R., Sheikh, S., Saxena, D., and Prokop, S. (2020). A patient-derived glioblastoma organoid model and biobank recapitulates inter-and intra-tumoral heterogeneity. Cell 180, 188–204. e122.

41. Milone, M.C., Fish, J.D., Carpenito, C., Carroll, R.G., Binder, G.K., Teachey, D., Samanta, M., Lakhal, M., Gloss, B., and Danet-Desnoyers, G. (2009). Chimeric receptors containing CD137 signal transduction domains mediate enhanced survival of T cells and increased antileukemic efficacy in vivo. Molecular therapy 17, 1453–1464.

42. Richman, S.A., Nunez-Cruz, S., Moghimi, B., Li, L.Z., Gershenson, Z.T., Mourelatos, Z., Barrett, D.M., Grupp, S.A., and Milone, M.C. (2018). High-affinity GD2-specific CAR T cells induce fatal encephalitis in a preclinical neuroblastoma model. Cancer immunology research 6, 36–46.

43. Wang, H., Chen, H., Liu, S., Zhang, J., Lu, H., Somasundaram, R., Choi, R., Zhang, G., Ou, L., and Scholler, J. (2021). Costimulation of γδTCR and TLR7/8 promotes Vδ2 T-cell antitumor activity by modulating mTOR pathway and APC function. Journal for immunotherapy of cancer 9.

44. Liu, S., Zhang, G., Guo, J., Chen, X., Lei, J., Ze, K., Dong, L., Dai, X., Gao, Y., and Song, D. (2018). Loss of Phd2 cooperates with BRAF V600E to drive melanomagenesis. Nature communications 9, 5426.

45. Hu, B., Ren, J., Luo, Y., Keith, B., Young, R.M., Scholler, J., Zhao, Y., and June, C.H. (2017). Augmentation of antitumor immunity by human and mouse CAR T cells secreting IL-18. Cell reports 20, 3025–3033.

46. Bronte, V., Brandau, S., Chen, S.-H., Colombo, M.P., Frey, A.B., Greten, T.F., Mandruzzato, S., Murray, P.J., Ochoa, A., and Ostrand-Rosenberg, S. (2016). Recommendations for myeloid-derived suppressor cell nomenclature and characterization standards. Nature communications 7, 12150.

47. Chen, G., Huang, A.C., Zhang, W., Zhang, G., Wu, M., Xu, W., Yu, Z., Yang, J., Wang, B., and Sun, H. (2018). Exosomal PD-L1 contributes to immunosuppression and is associated with anti-PD-1 response. Nature 560, 382–386.

48. Sherman, B.T., Hao, M., Qiu, J., Jiao, X., Baseler, M.W., Lane, H.C., Imamichi, T., and Chang, W. (2022). DAVID: a web server for functional enrichment analysis and functional annotation of gene lists (2021 update). Nucleic acids research 50, W216-W221.

49. Huang, D.W., Sherman, B.T., and Lempicki, R.A. (2009). Systematic and integrative analysis of large gene lists using DAVID bioinformatics resources. Nature protocols 4, 44–57.

50. Kaas, Q., Ruiz, M., and Lefranc, M.P. (2004). IMGT/3Dstructure-DB and IMGT/StructuralQuery, a database and a tool for immunoglobulin, T cell receptor and MHC structural data. Nucleic acids research 32, D208–D210.

51. Ehrenmann, F., Kaas, Q., and Lefranc, M.-P. (2010). IMGT/3Dstructure-DB and IMGT/DomainGapAlign: a database and a tool for immunoglobulins or antibodies, T cell receptors, MHC, IgSF and MhcSF. Nucleic acids research 38, D301–D307.

52. Rodrigues, J.P., Teixeira, J.M., Trellet, M., and Bonvin, A.M. (2018). Pdb-tools: a swiss army knife for molecular structures. F1000Research *7*.

53. Mongkolsapaya, J., Grimes, J.M., Chen, N., Xu, X.-N., Stuart, D.I., Jones, E.Y., and Screaton, G.R. (1999). Structure of the TRAIL–DR5 complex reveals mechanisms conferring specificity in apoptotic initiation. Nature structural biology 6, 1048–1053.

54. Honorato, R.V., Koukos, P.I., Jiménez-García, B., Tsaregorodtsev, A., Verlato, M., Giachetti, A., Rosato, A., and Bonvin, A.M. (2021). Structural biology in the clouds: the WeNMR-EOSC ecosystem. Frontiers in molecular biosciences 8, 729513.

55. Van Zundert, G., Rodrigues, J., Trellet, M., Schmitz, C., Kastritis, P., Karaca, E., Melquiond, A., van Dijk, M., De Vries, S., and Bonvin, A. (2016). The HADDOCK2. 2 web server: user-friendly integrative modeling of biomolecular complexes. Journal of molecular biology 428, 720-725.

56. Ou, L., Wang, H., Liu, Q., Zhang, J., Lu, H., Luo, L., Shi, C., Lin, S., Dong, L., and Guo, Y. (2021). Dichotomous and stable gamma delta T-cell number and function in healthy individuals. Journal for Immunotherapy of Cancer 9.

